# Specific Disruption of Established *P. aeruginosa* Biofilms Using Polymer-Attacking Enzymes

**DOI:** 10.1101/598979

**Authors:** Kristin N. Kovach, Derek Fleming, Kendra P. Rumbaugh, Vernita D. Gordon

**Affiliations:** Department of Physics and Center for Nonlinear Dynamics, The University of Texas at Austin, Austin TX 78712; Department of Surgery, Texas Tech University Health Sciences Center, Lubbock TX 79430; Institute for Cellular and Molecular Biology, The University of Texas at Austin, Austin TX 78712

## Abstract

Biofilms are communities of bacteria embedded in an extracellular matrix of self-produced polymeric substances. This polymer matrix lends the bacteria protection against a wide array of chemical and mechanical stresses that they may experience in their environment, which might be a location in the human body in the case of a biofilm infection, or a surface immersed in fluid in an industrial setting. Breaking down the matrix network renders biofilms more susceptible to physical disruption and to treatments. Different species of bacteria, and different strains within the same species, produce different types of matrix polymers – this suggests that targeting specific polymers for disruption may be more effective than non-specific approaches to disrupting biofilm matrices. In this study, we treated *Pseudomonas aeruginosa* biofilms with enzymes that are specific to different matrix polymers. We used bulk rheology to measure the resulting alteration in biofilm mechanics, and scanning electron microscopy to visualize the alteration in the matrix network upon treatment. Different lab strains of *P. aeruginosa* form biofilms that can be dominated by one of three main extracellular polysaccharides: Psl, alginate, and Pel, which binds electrostatically to extracellular DNA in the matrix. We applied enzymes to biofilms dominated by different extracellular polysaccharides and found that, for biofilms grown *in vitro*, the effect of enzymatic treatment is maximized when the enzyme is specific to a dominant matrix polymer – for such a case, specifically-matched enzymatic treatment tends to: reduce yield strain and yield stress; reduce or eliminate long-range structure and shorten or eliminate connecting network fibers in the biofilm as seen under scanning electron microscopy; and increase the rate of biofilm drying, most likely due to increased diffusivity as a result of network compromise. However, for *ex vivo* biofilms grown in murine wounds, we find that generic glycoside hydrolases have more profound disruptive effects than specifically-matched enzymes, even though they had no measurable effect for biofilms grown *in vitro*. This highlights the importance of the environment in which the biofilms are grown, the need to take this into account when developing treatments for biofilms, and the possibility that effective approaches to eradicating biofilms in environmental or industrial settings may need to be very different from effective treatments of infection.

## Introduction

*Pseudomonas aeruginosa* is an opportunistic human pathogen, known for its ability to form robust biofilms. Bacterial biofilms consist of bacteria in a complex matrix of extracellular polymers and proteins. Multiple types of matrix polymers can be produced, and the importance of a specific polymer type varies with biofilm-forming species and strain. For biofilm infections, the matrix protects the inhabitant bacteria chemically by inhibiting the diffusion of antibiotics into the biofilm and by binding to antibacterial chemicals produced by the host immune response.^1–3^ Furthermore, the mechanical integrity and structure of the bacterial biofilm conferred by the matrix gives rise to stable microenvironments that contribute to phenotypic antibiotic tolerance.^4–5^ Thus, both the chemical and the structural properties of the biofilm matrix contribute to bacterial tolerance of harsh environments.

Many common interventions in *P. aeruginosa* infections, like antibiotic treatment, are far more successful if the bacteria are in a planktonic state rather than in a biofilm.^6^ It has been shown that dispersing biofilms, or breaking them up into smaller pieces, can increase the efficacy of antibiotics in killing or inhibiting bacteria, as well as increasing the ability of phagocytic immune cells to engulf and kill bacteria.^6–9^ However, in cases such that the patient already has a well-established biofilm infection, finding methods to mechanically compromise and disrupt biofilms without harming the patient is often non-trivial. *P. aeruginosa* biofilms have proven mechanically resilient against many simple perturbations such as ionic disruption, pH changes, and addition of small organic molecules, hinting that once the matrix network has formed, connections are difficult to break.^10^ Therefore, more and better methods for disrupting established biofilms are needed. The glycoside hydrolases cellulase and α–amylase have successfully been used against *P. aeruginosa* biofilms, disrupting the biofilms and rendering them more susceptible to antibiotics *in vivo* and *in vitro*.^8–9^ However, these enzymes are not specific against any components of the *P. aeruginosa* biofilm matrix, and we might expect more specific enzymes to have greater efficiency than generic glycoside hydrolases, with less potential for harming the host. That being said, biofilms formed by different strains of *P. aeruginosa* are dominated by different types of extracellular polymers (EPS) and will therefore respond differently to enzymes that attack specific polymers. Therefore, it is important to test both specific and non-specific enzymes against these biofilm infections in order to determine the best approach to resolving them.

The primary EPS constituents of *P. aeruginosa* biofilm matrices are the polysaccharides alginate, Psl, Pel, and extracellular DNA.^11–12^ The enzyme alginate lyase has successfully been used to disperse biofilms, rendering them more susceptible to the immune system and antibiotics.^7, 13–16^ DNase is another enzyme that is successful in dispersing and rendering biofilms susceptible to antibiotics.^17–18^ Psl-and Pel-specific glycoside hydrolases have been synthesized to attack *P. aeruginosa* biofilms and they increase antibiotic susceptibility when they are used to treat Psl-and Pel-dominant biofilms, respectively. In addition, these hydrolases were able to decrease biomass in a wide variety of *P. aeruginosa* strains to varying effectiveness, depending on the hydrolase and strain.^19^

In this study, we grew biofilms *in vitro* from lab strains of *P. aeruginosa* with different dominant extracellular polysaccharides. Once the mature biofilms were well-established, we treated them with enzymes that are specific to different matrix polymers. We used bulk oscillatory rheology to quantify the changes in biofilm mechanics resulting from enzymatic treatment. Alginate lyase has a bigger effect on the mechanics of alginate-dominant biofilms than Psl-dominant or Pel-dominant biofilms. DNase I changes the mechanics of Pel-dominant biofilms more than Psl-dominant or alginate-dominant biofilms; Pel has been shown to bind to extracellular DNA (eDNA) in the matrix.^20^ Thus, we found that the largest changes in biofilm mechanics occur when the enzyme is matched to a dominant polymer in the biofilm matrix. Biofilms that have had their mechanics compromised by specific enzymatic treatment also have a reduced ability to hold water and stay hydrated; we attribute this to the effects of the compromised network mesh on the increased diffusive transport of water out of the biofilm. Upon treatment with the glycoside hydrolases cellulase and α-amylase, which are not specific to matrix polymers in the *in vitro* biofilms we study, the mechanical alterations are small and not significant. However, our findings for biofilms grown *in vitro* do not extend to the dispersal of biofilms grown *in vivo*, in a mouse model of wound infection, and then excised and treated *ex vivo*. For *ex vivo* biofilms, the generic glycoside hydrolases are more effective than the specific enzymes at inducing dispersal regardless of the lab strain of bacteria used. The mis-match between mechanical changes for biofilms grown *in vitro* and dispersal of biofilms grown *in vivo* could result if the matrix composition of biofilms grown *in vivo* is significantly different from that of biofilms grown *in vitro* by genetically-identical strains of bacteria, and thus highlights the importance of the biofilm growth environment and the need for taking growth conditions into account when devising anti-biofilm strategies. Alternatively or in addition, the mis-match between our findings for *in vitro*-and *in vivo*-grown biofilms may indicate that the dispersal of constituent bacteria does not depend primarily on the mechanical integrity, or lack thereof, of the embedding biofilm matrix.

## Background

### P. aeruginosa matrix polymers and their effects on biofilm mechanics

Our previous work and the work of others have shown that increased production of Pel or alginate will increase the yield strain of the biofilm; yield strain measures how large of a deformation can be sustained before material failure begins.^21–23^ Alginate is a negatively-charged polymer, and increased alginate production increases yield strain while decreasing the elastic modulus.

Pel is a positively-charged chain of amino sugars, and is thought to bind ionically with extracellular DNA (eDNA) in the biofilm matrix; eDNA incorporated into the biofilm matrix can come from lysed bacteria or from non-biofilm cells, including host immune cells.^24–26^ Linking two polymers (eDNA and Pel) may be what allows high Pel production to increase the yield strain of biofilms without reducing elastic modulus. A possible analogue may be found in synthetic hydrogels, in which two polymers can be made to complex with one another such that one polymer acts as a brittle “sacrificial” bond so that when sheared, the gel will fracture at these bonds, allowing the second polymer to extend to allow large deformations.^27^ These so-called double network gels are very tough against mechanical perturbations.^28^

Psl is a neutral, branched polymer, and increasing Psl production increases the elastic modulus of the biofilm as long as the Psl-binding protein CdrA is also produced.^21^ Combining high yield strain and high elasticity can make a strong polymer matrix, resistant to a variety of physical stresses. Indeed, it seems that *P. aeruginosa* may often evolve to this end. For example, in chronic Cystic Fibrosis (CF) infections, that can last as long as decades, the production of both Psl and alginate are increased as the infecting population evolves inside the patient.^29^

### Enzymes targeting specific matrix polymers

Enzymes target chemically-specific substrates, although some are more discriminating than others. Alginate is made up of monomer units of α-L-guluronate and β-D-mannuronate. The order and repetition of these units is highly dependent upon the source of alginate, with variation also present in single alginate sources.^30^ Alginate lyase targets the glycosidic linkages in the alginate polysaccharide, breaking it down into smaller oligosaccharides via a β-elimination reaction at β-1,4 bonds.^30^ Alginate lyase has been well studied as a treatment mechanism for *P. aeruginosa* biofilms, as alginate is known to provide many chemical and physical protections for bacteria. These studies have found that alginate lyase can degrade biofilms, increase antibiotic susceptibility, enhance functionality of the host immune response, and alter the viscosity of sputum in patients with CF.^7, 13, 16, 31-33^

DNase I cleaves phosphodiester bonds in the backbone of DNA via hydrolysis, breaking it down into pieces as small as 10 base pairs. Like alginate lyase, DNase I is also thought to be a promising enzyme for treatment of *P. aeruginosa* biofilms, since extracellular DNA can be an important component of *P. aeruginosa* bacterial biofilms, lending structural integrity and contributing to antibiotic resistance.^12^ DNase I is already a common treatment for patients with CF^34^ and, much like alginate lyase, has proven successful in degrading biofilms, increasing antibiotic susceptibility, and altering the viscosity of CF sputum.^12, 17-18, 35^

The glycoside hydrolases cellulase and α-amylase both break down polysaccharides by hydrolysis of glycosidic linkages. Cellulase hydrolyzes β-1,4 bonds and α-amylase hydrolyzes α-1,4 bonds in polysaccharide chains. While cellulase and α-amylase specifically decompose cellulose and amylose, respectively, they have also shown the ability to break down other polysaccharides with structural similarities to their namesakes.^9^ Both cellulase and α-amylase can inhibit *P. aeruginosa* biofilm growth, and they have been shown to degrade biofilms and render bacteria more susceptible to antibiotics in wounds.^8-9, 36-37^

## Results

Enzymatic treatments were applied in parallel with control treatments. To minimize variations in culturing conditions such as humidity on the day of growth, the biofilms for each pair of treatment and control were initiated from the same overnight liquid culture and grown in the same incubator, on nutrient agar plates from the same preparation batch. For each pair of treatment and control biofilm, rheological measurements were performed on the same day, in immediate succession. To determine the effects of the treatments on different mechanical properties, we report a ratio of the value of a mechanical property (elastic modulus, yield strain, or yield stress) for a treated biofilm to the value of the same property for the corresponding control biofilm. This is very similar to the approach we took in our earlier work, and is intended to account for the effects of day-to-day variation in the measured mechanics of biofilms grown from the same bacterial strain.^21^

To study the response of individual polymers to each form of disruption, we use four PAO1 lab strains of *P. aeruginosa*, three of which have been genetically modified to alter polysaccharide production. *In vitro*, the wild type (WT) of strain PAO1 produces primarily Pel and Psl, with alginate having no mechanical contribution to the biofilm.^38^ To isolate the behavior of Pel and Psl, we use the strains Δ*wspF*Δ*psl* and Δ*wspF*Δ*pel*, respectively. The deletion of the *wspF* gene causes overproduction of cyclic-di-GMP, an intracellular signaling molecule that increases the constitutive expression of both Pel and Psl in the biofilm.^39–40^ Deleting either the *pel* or *psl* gene in addition to *wspF* forces the biofilm to overproduce the remaining polysaccharide. In the figures, Δ*wspF*Δ*psl* is denoted as Pel+, and Δ*wspF*Δ*pel* is denoted as Psl+. To learn about the contribution of alginate to the biofilm, we used PAO1 Δ*mucA*.^41^ The *mucA* gene regulates the production of alginate, and so by deleting this gene, alginate is overproduced. Although Pel and Psl are present in the Δ*mucA* biofilm, we have previously shown that alginate has a strong influence on mechanical properties for Δ*mucA* biofilms.^21^ In the figures, Δ*mucA* is denoted as Alg+.

### Alginate lyase and DNase I act on biofilm matrices with specificity

We find that treating Psl+, Pel+, and Alg+ biofilms with 200 U/mL alginate lyase causes statistically-significant changes in the mechanical properties of both Pel+ and Alg+ biofilms. The effect on the Alg+ biofilm is greater than the effect on the Pel+ biofilm, as the elastic modulus and the yield stress of the Alg+ biofilm both increase by more than a factor of five. In contrast, the elastic modulus of the Pel+ biofilm increases by a factor of less than 1.5, and its yield stress actually decreases due to the treatment causing a decrease in the yield strain (Figure 1).

**Figure 1.**
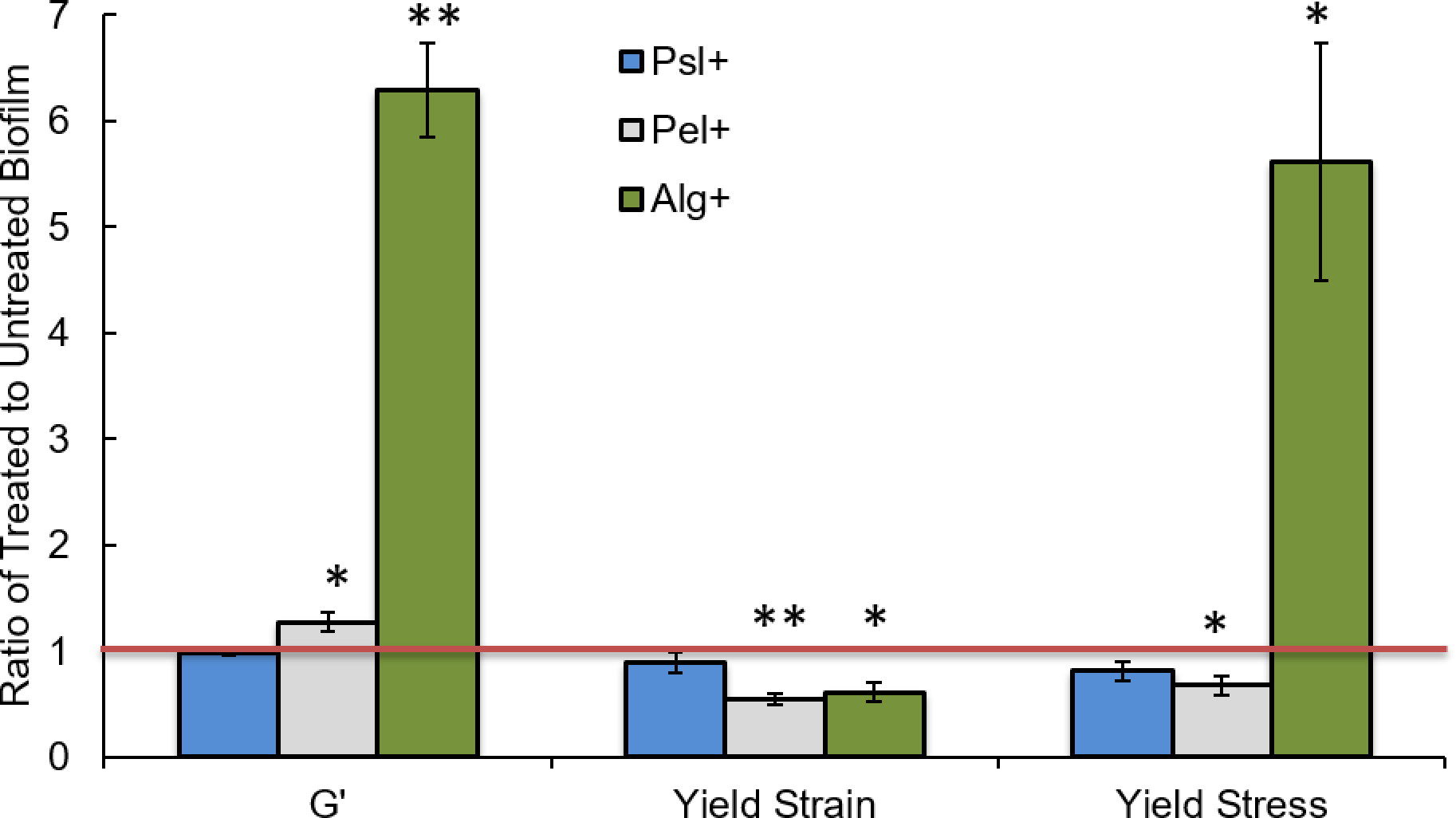
Treatment by 200 U/mL alginate lyase impacts the mechanics of alginate-dominant biofilms and Pel-dominant biofilms. The mechanical properties measured for each treated biofilm are compared to the corresponding property measured for a biofilm treated by the control solution by taking ratios. Thus, a ratio of one indicates no change upon treatment with the enzyme, a ratio greater than one indicates an increase of that mechanical property upon treatment, and a ratio less than one indicates a decrease in that mechanical property upon treatment. For alginate-dominant biofilms, there is both a 6.3x increase in elastic modulus and a 1.6x decrease in yield strain. For Pel-dominant biofilms, there is a 1.3x increase in elastic modulus and a 2x decrease in yield strain. * p ≤0.05 and ** p≤0.01. Error bars are standard error of the mean.

In previous studies by others, the dispersal properties of alginate lyase were found to be catalysis-independent; merely the presence of any protein triggered a generic dispersal signal in *P. aeruginosa*.^31^ However, the strongly specific activity of alginate lyase on Alg+ biofilms that we measure in our experiments indicates that the enzyme is specifically cleaving the alginate. As a further check to confirm the importance of enzymatic specificity in this case, we treat an Alg+ biofilm with arginine, a main amino acid component of alginate lyase, to see if our rheological measurements are sensitive to the protein-triggered dispersal previously seen to result from the addition of arginine.^31^ Treatment with arginine had no effect on the Alg+ biofilm (Figure S1). This is additional evidence that the changes we measure in biofilm mechanics are specific to the effect of alginate lyase on alginate in the matrix. Treatment with arginine had some effect on the WT biofilm—with matrix composed primarily of Psl and secondarily of Pel—but the mechanical alteration did not mirror that of alginate lyase (Figure S2).

Despite the effect of alginate lyase on the Pel+ biofilms, it seems unlikely that alginate lyase is catalytically active on the Pel polysaccharide. Pel and alginate are composed of different monomer units; Pel is composed of N-acetylglucosamine and N-acetylgalactosamine.^24^ They also have opposite charges, with Pel being cationic and alginate being anionic. While the Pel+ biofilm should only have minimal alginate polysaccharide present, it is possible that the small amount present interacts electrostatically with the Pel polysaccharide network in such a way as to increase yield strain, as we have previously suggested may happen for anionic eDNA binding electrostatically to Pel in the biofilm matrix. We address why cleaving matrix polymers results in a higher measured elastic modulus below, in the subsection “*The increase in elastic modulus is an effect of drying*.”

When biofilms are treated with 500 U/mL DNase I, we find that the elastic modulus of Pel+ biofilms, and no other biofilm type, increases by more than a factor of two (Figure 2). This parallels an effect of alginate lyase on Alg+ biofilms. Pel is thought to associate with eDNA in the matrix^20^, and our mechanical measurements are consistent with the idea that Pel and eDNA interact to mechanically stabilize these biofilms.

**Figure 2.**
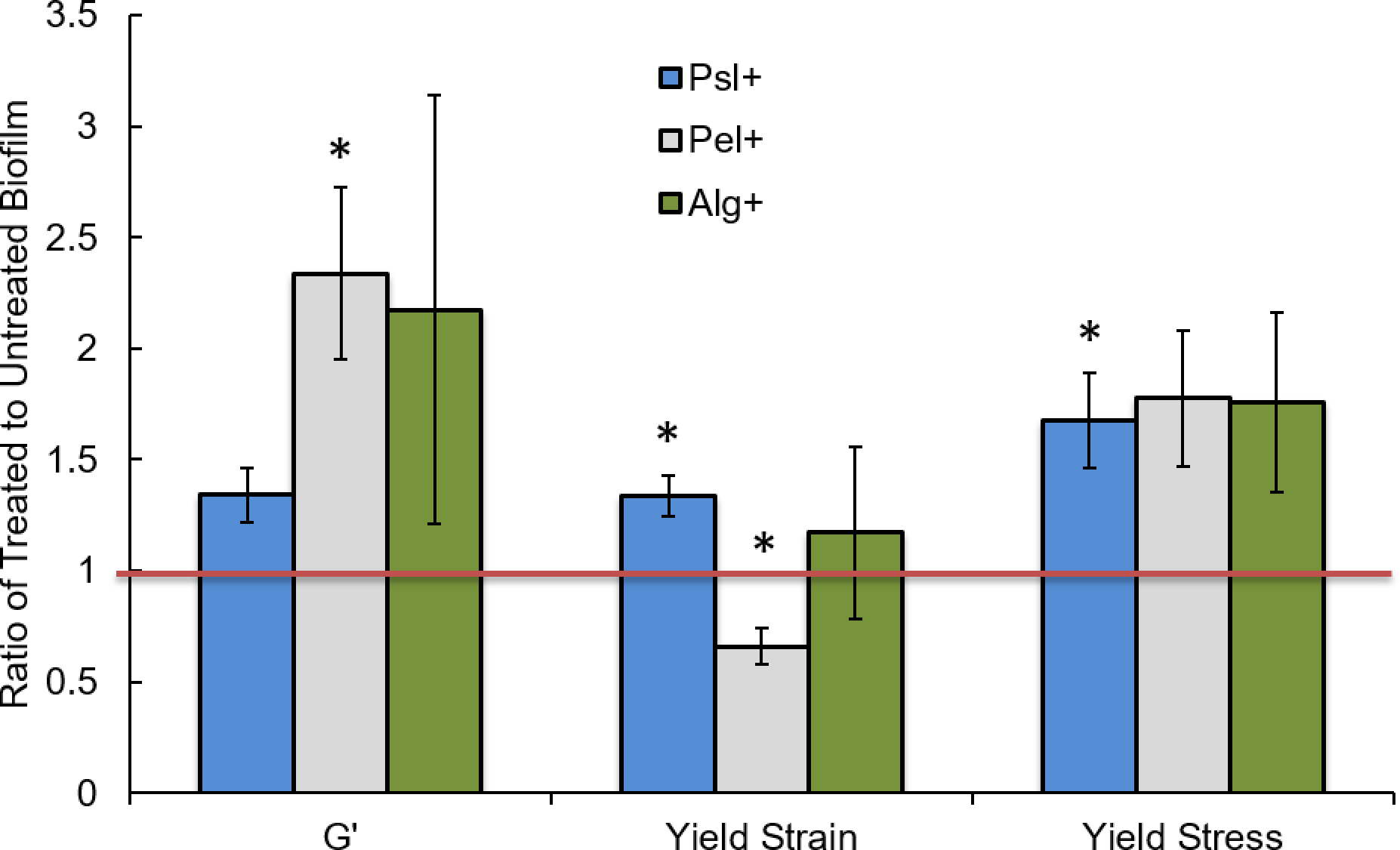
Treatment by 500 U/mL DNase I impacts the mechanics of Pel-dominant biofilms only. The mechanical properties measured for each treated biofilm are compared to the corresponding property measured for a biofilm treated by the control solution by taking ratios. Thus, a ratio of one indicates no change upon treatment with the enzyme, a ratio greater than one indicates an increase of that mechanical property upon treatment, and a ratio less than one indicates a decrease in that mechanical property upon treatment. For Pel-dominant biofilms, there is a 2.3x increase in elastic modulus and a 1.6x decrease in yield strain. * p ≤0.05 and ** p≤0.01. Error bars are standard error of the mean.

DNase I has no statistically-significant impact on Alg+ biofilms. However, DNase I does slightly increase the yield strain and yield stress of Psl+ biofilms. The mechanism underlying this result is less clear, but it likely reflects the smaller, but still present, role of eDNA as a structural constituent in these biofilms.^12, 42-43^

### Alginate lyase and DNase I decrease yield strain

When 200U/mL alginate lyase acts on Alg+ and Pel+ biofilms and when 500 U/mL DNase I acts on Pel+ biofilms, the yield strain decreases. Alginate lyase decreases the yield strain of Alg+ biofilms by ~40% and that of Pel+ biofilms by ~45% (Figure 1). DNase I decreases the yield strain of Pel+ biofilms by ~35% (Figure 2). The decrease in yield strain likely occurs due to the chains of the polymer network being shortened, so that that the matrix cannot maintain integrity in the face of large deformations.

### Mechanical alterations are dose-dependent

To probe how mechanical changes depend on the amount of polymer cleavage, we vary the concentration of the enzyme treatments. At lower enzyme concentrations, the only statistically significant mechanical change is a ~130% increase in elastic modulus when we treat an Alg+ biofilm with 20 U/mL alginate lyase, compared to the ~530% increase in elastic modulus when we treat with 200 U/mL alginate lyase (Figure 3). Similarly for DNase I, treating Pel+ biofilms with an enzyme concentration of 500 U/mL has statistically-significant effects, but a concentration of 50 U/mL does not (Figure 4).

**Figure 3.**
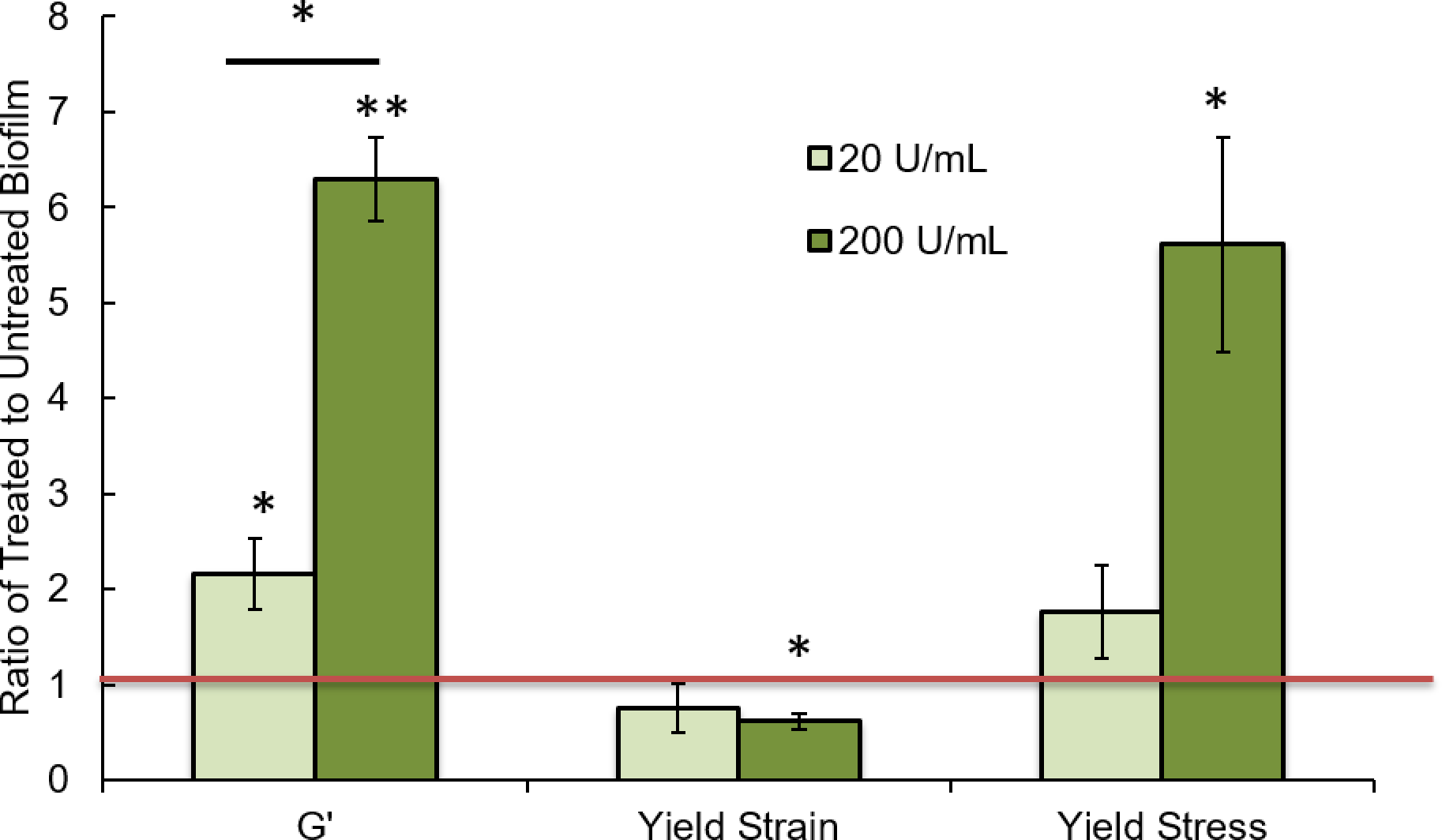
The effects of alginate lyase treatment on alginate-dominant biofilms depends on dosage. When alginate lyase treatment increases from 20 U/mL to 200 U/mL, the increase in elastic modulus goes from 2.2x to 6.3x. The change in yield strain goes from being non-significant to a 1.6x decrease.

**Figure 4.**
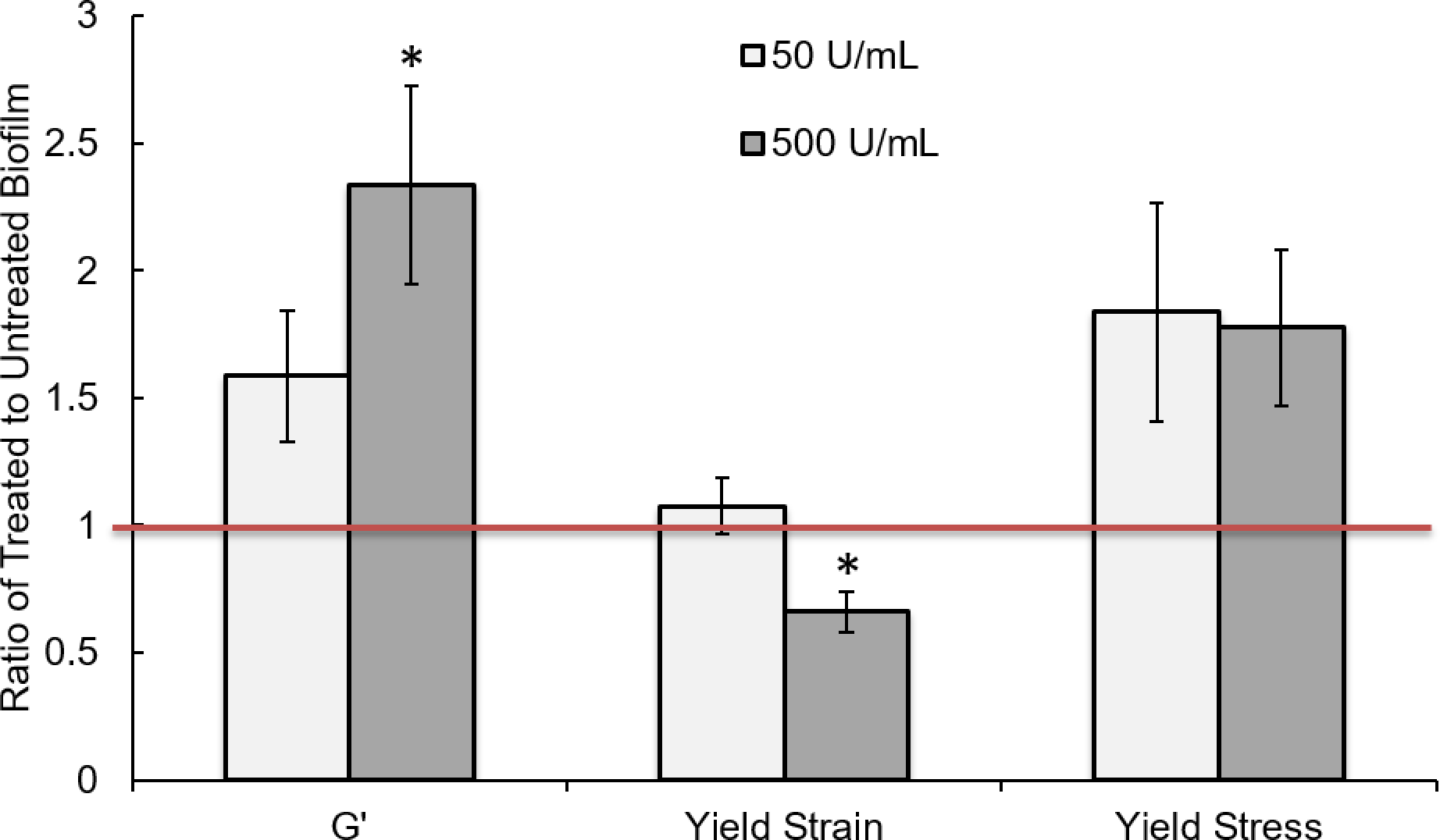
The effects of DNase I treatment on Pel-dominant biofilms depends on dosage. When DNase treatment increases from 50 U/mL to 500 U/mL, the effects of the treatment on elastic modulus and yield strain become statistically significant.

### Electron microscopy indicates specific disruption of matrix structure

To visualize network compromise, we use scanning electron microscopy (SEM) to image Alg+ and Psl+ biofilms both untreated and treated with alginate lyase (Figures S3 and S5) and Pel+ and Psl+ biofilms both untreated and treated with DNase I (Figures 5 and S4). Most dramatically, we see that an untreated Pel+ biofilm has 100-micron scale structures (Figure 5C) and that the cells are embedded in an interconnected network of stringy polysaccharides (Figure 5D). When treated with DNase I, these large structures and surface attachment of cells are compromised (Figure 5A), and there are less network strands present (Figure 5B). It appears that the loss of yield strain in Pel+ biofilms due to treatment by DNase I is most likely due to the reliance of the Pel network on extracellular DNA for its ability to yield without breaking.

**Figure 5.**
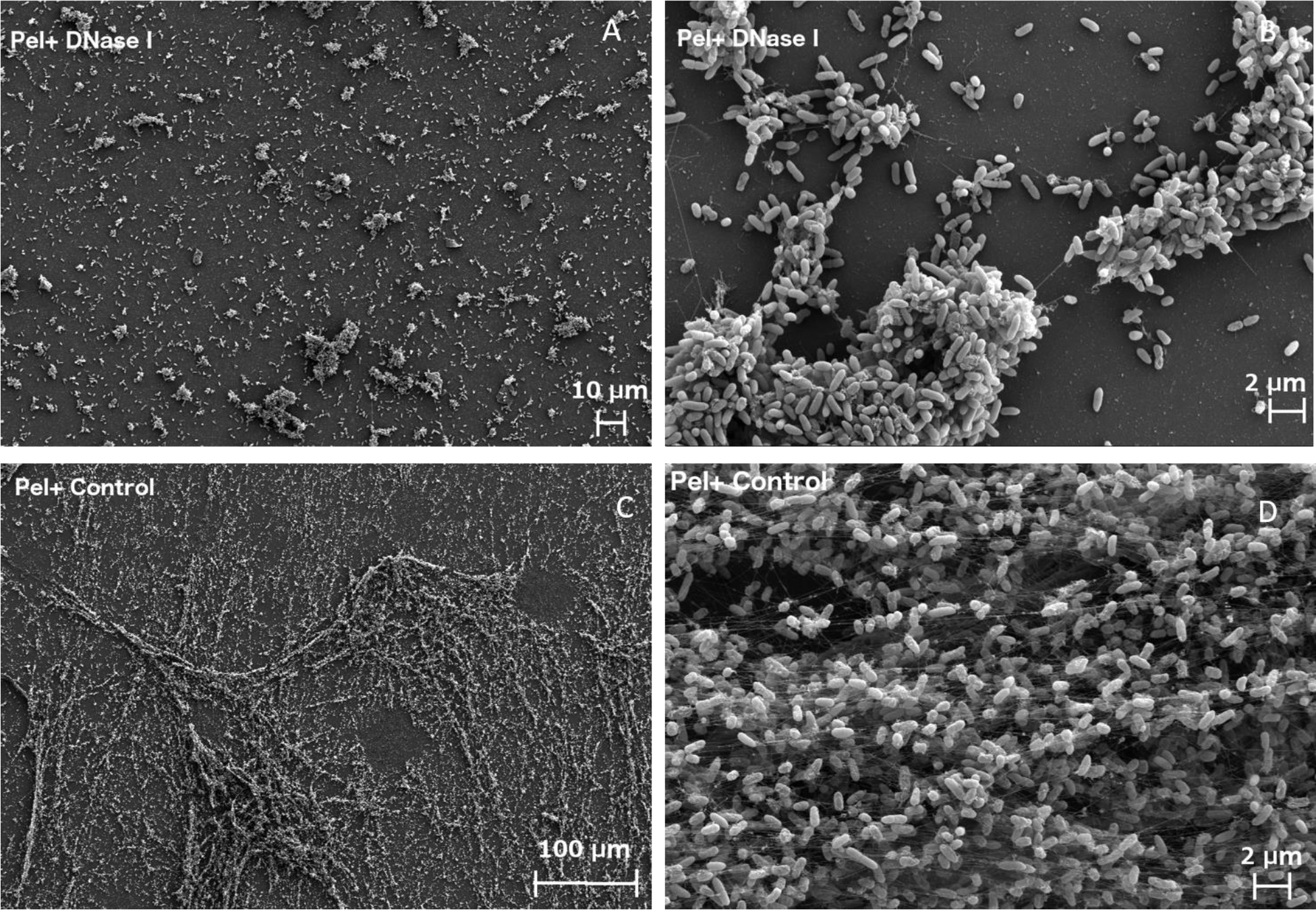
SEM images of Pel-dominant biofilms. (A, B) Biofilms that have been treated by DNase I show (A) no long-range interconnectivity and (B) no polymer fiber strands. In contrast, control biofilms that were not treated by DNAse I have (C) long-range interconnectivity and structure and (B) visible polymer fibers bridging bacteria.

Despite alginate lyase causing mechanical changes in Alg+ biofilms, there is not a dramatic difference in the SEM images of treated and untreated biofilm (Figure S3). In our Alg+ biofilms, Pel and Psl polysaccharides are still present, and so it may be that the alteration in Alg+ polysaccharide is not visible in SEM imaging. An important consideration is that the preparation steps for SEM cause violent agitation to the biofilm formations on the surface. As such, the SEM preparation process most likely selects for the most strongly surface-attached cellular structures. If cells attached to the surface and one another via alginate have impaired surface attachment, we may only be imaging cells that were attached to the surface with Pel or Psl polysaccharide once SEM preparation is complete. In addition, alginate-overproducing cells tend to make less cohesive biofilms even without any treatment, and the fixation process likely removes the weakest of polysaccharide connections—which in the case of the alginate-overproducing cells are relatively weak in both treated and untreated samples. Indeed, it has been shown with simulations of enzymatic treatment on biofilms that if the EPS network does not contribute to cohesiveness, surface growth is difficult to remove.^44^ As such, an alginate biofilm may be difficult to alter at the surface.

We also imaged the effects of enzyme treatments on Psl+ biofilm, for which no mechanical compromise occurred. There appears to be some alteration in surface cell-density in treatment with DNase I on Psl+ biofilm (Figure S4). We speculate that, although extracellular DNA does not play a major role in the mechanical properties of Psl+ biofilms, it does still play a role in cellular attachment to the surface. Alginate lyase also appears to slightly affect Psl+ biofilm surface attachment (Figure S5A and S5C) as well as alter the visible polysaccharide network (Figure S5B and S5D). As there may be small amounts of alginate present in our Psl+ biofilms, it may be that these alginate polysaccharides do play some role in surface attachment, while not being mechanically important to the bulk matrix. Surprisingly, there are dense structures present surrounding cells in Psl+ biofilms treated with both of these enzymes (Figure S4B and S5B). One possible explanation for these structures is biofilm dehydration due to enzyme treatment, as discussed in the next section. If the polysaccharide network collapses on itself due to lack of hydration, the network might appear as dense structures in our SEM images.

### The increase in elastic modulus is an effect of drying

Unlike the decrease in yield strain caused by cleaving matrix polymers, it is not immediately obvious why enzymatic treatment, resulting in shorter polymers, should increase elastic modulus. Of greater pragmatic concern, an increase in elastic modulus (and therefore potentially an increase in yield stress as well), would seem likely to be protective for the bacteria in the biofilm, and therefore not a desirable outcome if the specific enzymes alginate lyase and DNase are to be considered as an approach to clinical treatment.

However, the interpretation of these *in vitro* experiments is more complex than it may at first appear. A modified version of Fick’s first law of diffusion gives us the drying of a polymer gel or network related to the diffusion coefficient of the solvent by

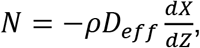

where N is the drying rate, ρ is the volume density of polymer, D_eff_ is the effective diffusion coefficient of the solvent, X is the solvent content, and Z is the thickness of the matrix.^45^ By cleaving polymer chains, enzymes reduce the connectivity of the biofilm matrix and therefore allow water to diffuse more freely through the matrix.

In order to verify that the biofilms are drying, biofilms are weighed before and after treatment with enzymes and a control. When an Alg+ biofilm is treated with alginate lyase, it dries ~50% more than a biofilm treated with a control (Figure 6). When a Pel+ biofilm is treated with DNase I, it dries ~20% more than the control (Figure 6). In addition, by visual inspection, we see that the well-hydrated mucoid biofilm does indeed appear to have lost much of its water content (Figure 7).

**Figure 6.**
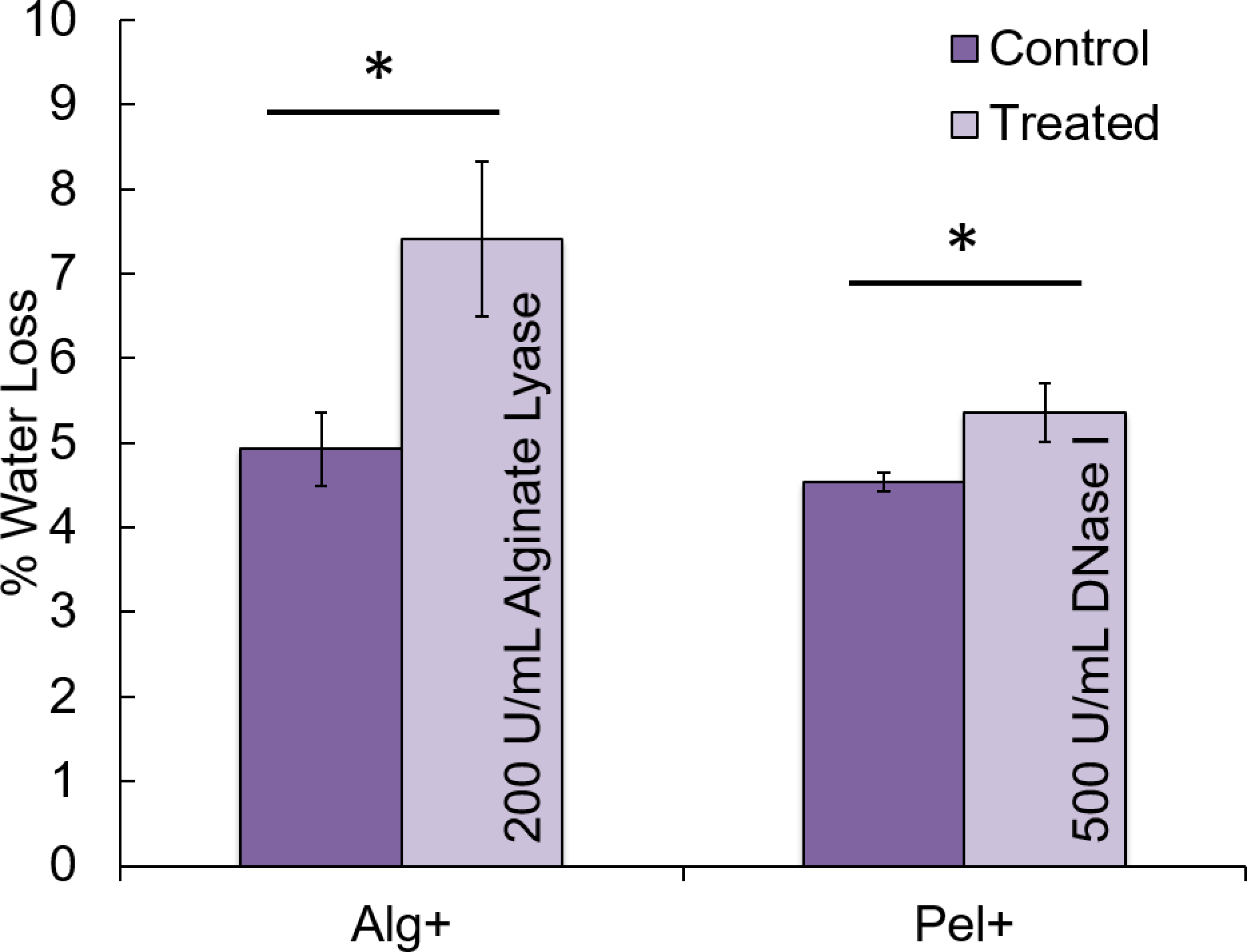
Enzymatically-disrupted biofilms dry more quickly than undisrupted biofilms. When alginate-dominant biofilms are treated with 200 U/mL alginate lyase, on average they lose 7.4% of their weight to water loss, while the control biofilms lose only 5%. When pel-dominant biofilms are treated with DNase I, on average they lose 4.5% of their weight to water loss, while the control biofilms lose only 5.4%.

**Figure 7.**
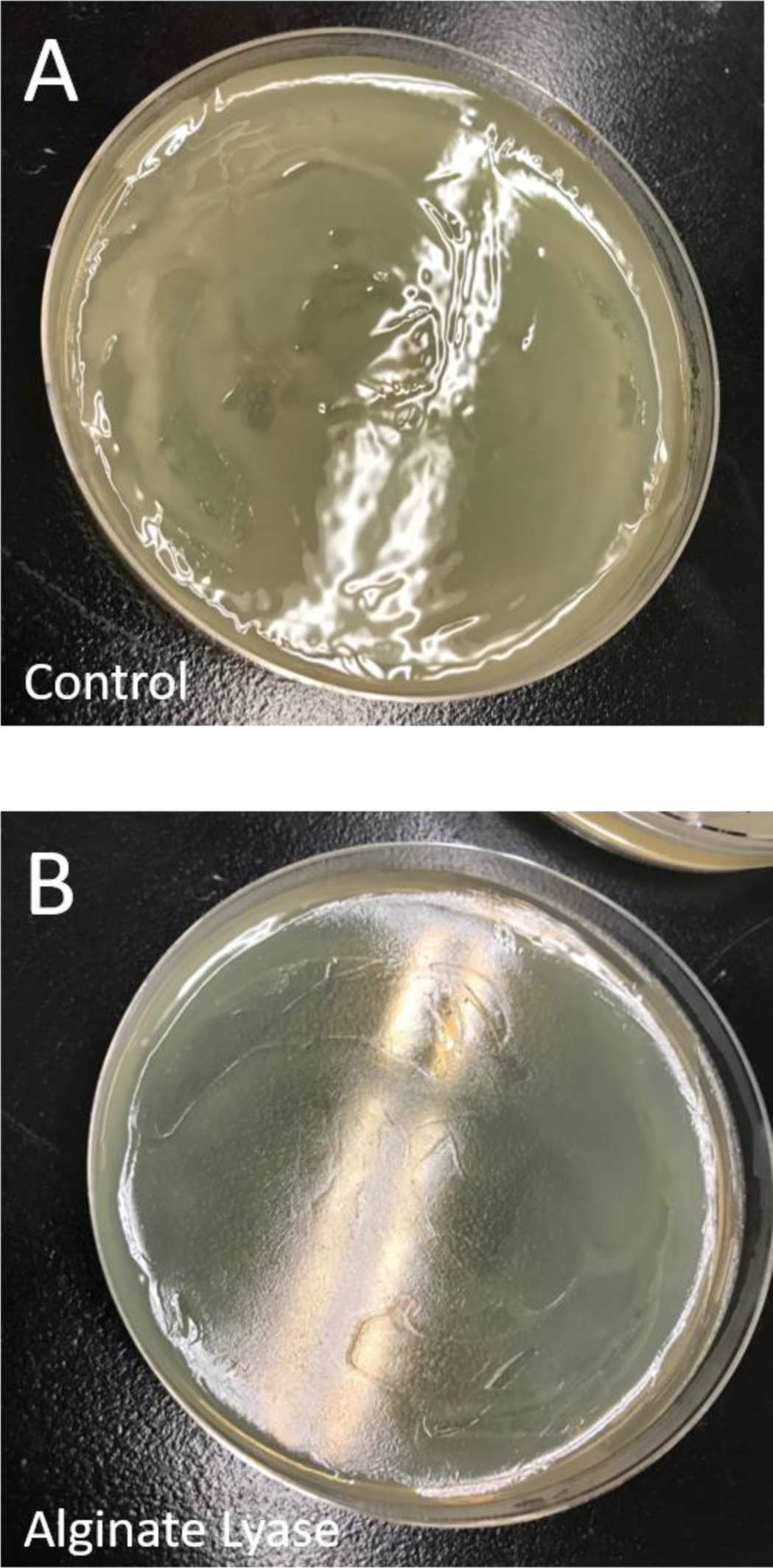
A) Alginate-dominant biofilm treated by control appears wetter than B) an alginate-dominant biofilm treated by 200 U/mL alginate lyase, which appears dry.

Thus, during the hour of enzyme treatment at 37°C, enzyme-treated biofilms dry more quickly than do biofilms treated with an inactive control. Therefore, when biofilms are measured in the rheometer, the enzymatically-compromised biofilms contain less water and therefore have a higher polymer concentration than do the corresponding control biofilms. It is well known that elastic modulus scales with polymer density G’∝ c^A^, where c is polymer concentration and A is a scaling factor (A=2.25 for an entangled polymer in a good solvent).^46^ Thus, the increase in elastic modulus most likely reveals a marked drop in polymer moisture content due to the polymer losing the ability to hold water, rather than any increase in structural cohesiveness or integrity.

When an Alg+ biofilm is treated with 200 U/mL alginate lyase, we see a dramatic ~530% increase in elastic modulus. 200 U/mL alginate lyase also causes a ~30% increase in elastic modulus of Pel+ biofilms (Figure 1). As with yield strain, we see that alginate lyase acts on Alg+ and Pel+ biofilms, although in this case to different magnitudes. Thus, while it seems that alginate lyase is causing a larger change in response for Alg+ than Pel+, it may simply be that Alg+ biofilms hold more water natively, so when the matrix is compromised, there is a more dramatic increase in polymer density as the matrix loses water. If this is true, 200 U/mL seems to affect Alg+ and Pel+ the same with respect to matrix compromise.

When 500 U/mL DNase I acts on Pel+ biofilms, the elastic modulus increases by ~130% (Figure 2). As such, it seems that DNase I can also cause drying in the polymer matrix due to the breakdown of matrix integrity. An alternative explanation for the measured increases in elastic modulus upon enzymatic treatment would be if new cross-link points form at the ends of the cut chains, thereby increasing cross-link density. However, if the biofilms had potential for new cross-link points, we would expect that exogenously added extracellular DNA would be incorporated into the already-formed matrix. We see no significant mechanical changes in Pel+ biofilms when eDNA is added after biofilm growth (Figure 8). Although it may be that the exogenous eDNA units cannot replace whatever links the biofilm eDNA has already formed, this also indicates that there is not an excess of cross-link points to reinforce a matrix of shorter DNA chains in the case of DNase I addition. From this, we can presume that there is matrix compromise with the addition of DNase I, with no new cross-link points, and that the rise in elastic modulus is a result of increased water diffusion and subsequent drying.

**Figure 8.**
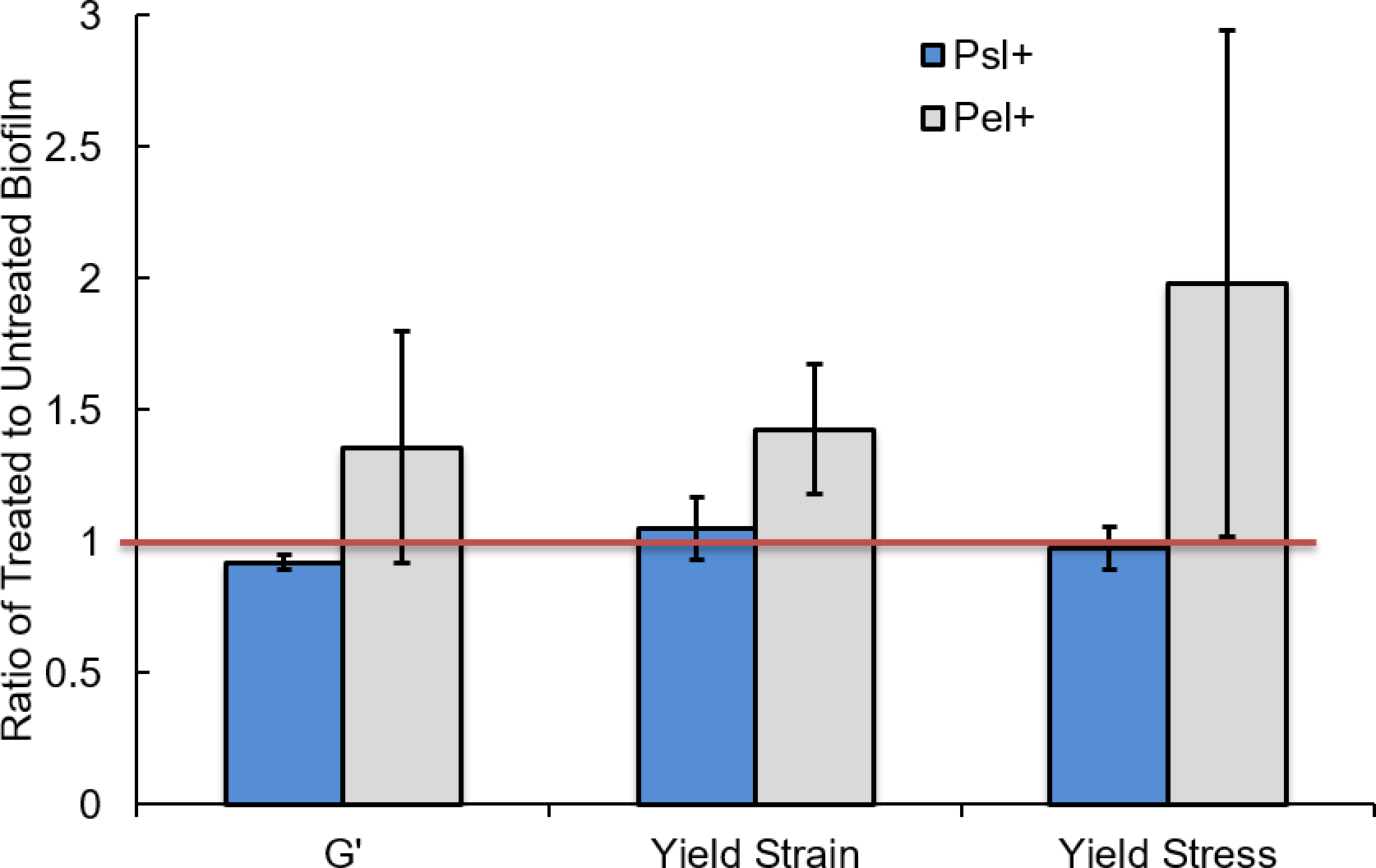
The addition of exogenous extracellular DNA after biofilms have grown does not affect the mechanical properties of biofilms. After the biofilms were grown, ~7mg/mL of eDNA were added to each, Each piece of eDNA was ≤ 2000 base pairs. * p ≤0.05 and ** p≤0.01. Error bars are standard error of the mean.

In summary, alginate lyase and DNase I are very different enzymes with entirely different substrate specificity, and yet, when they compromise a biofilm that is dominated by a polymer matched to their specific activity, the resulting mechanical properties are similar: a decrease in yield strain and, when the environmental conditions are such that drying is possible, an increase in elastic modulus. It is important to note that for biofilms infecting the body, or for any other condition where there is surrounding fluid or high humidity, increased diffusion would **<u>not</u>** be expected to result in drying although transport of antibiotics, immune factors, and other chemicals would be increased.

### Glycoside hydrolases cellulase and α-amylase do not alter mechanics of any biofilm type tested

Our results above suggest that the efficacy of enzymes in altering biofilm mechanics is likely to strongly depend on enzyme-specific activity against a dominant matrix constituent. However, others have found that cellulase and α-amylase, which are not specific to any biofilm matrix component, are successful in inhibiting and dispersing *P. aeruginosa* biofilms.^9^ Therefore, we tested the effect of both cellulase and α-amylase on the mechanics of Psl+, Pel+, and Alg+ biofilms.

Upon treatment with 5% α-amylase, Psl+ biofilms experience a ~5% decrease in elastic modulus, and Alg+ biofilms experience a ~40% increase in yield strain; no other statistically-significant effects are seen (Figure 9). Upon treatment with 5% cellulase, the yield stress of Alg+ biofilms decreases by ~60%; no other statistically-significant effects are seen (Figure 10). Thus, compared with the more than five-fold increase in elastic modulus and yield stress experienced by Alg+ biofilms upon treatment with alginate lyase, and the more than two-fold increase in elastic modulus experienced by Pel+ biofilms upon treatment with DNase I, the effects of non-specific glycoside hydrolases on biofilm mechanics are negligibly small. It is notable that the increase in yield strain experienced by Alg+ biofilms treated by α-amylase parallels, for our results with specific enzymes, only the increase in yield strain seen for Psl+ biofilms treated by DNase I, and that we do not expect specific activity of DNase I against the Psl extracellular polysaccharide.

**Figure 9.**
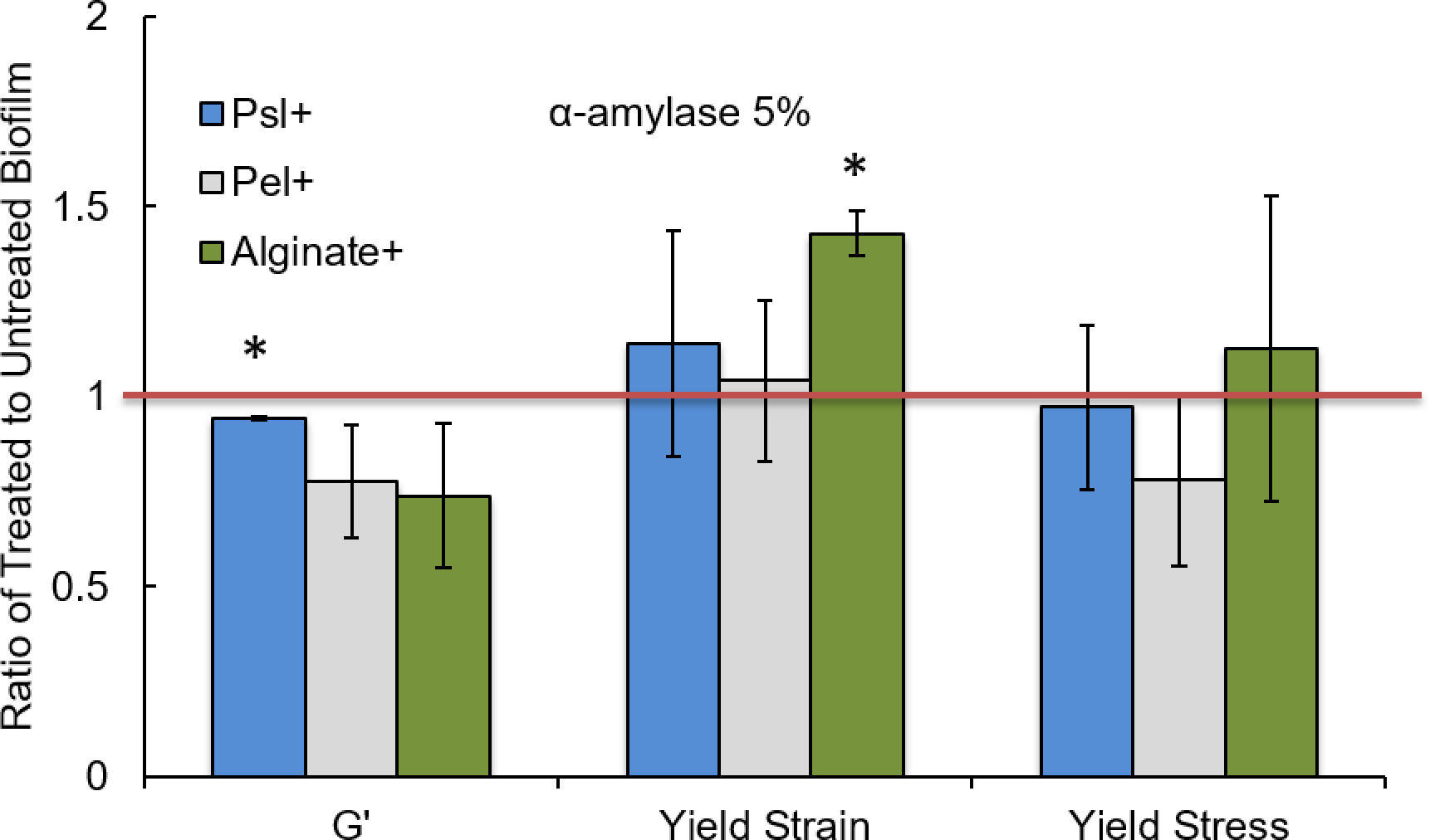
Treatment by 5% α-amylase has minimal effects on biofilm mechanical properties. Psl-dominant biofilms experience a slight decrease in elastic modulus while alginate-dominant biofilms have an increase in yield strain. * p ≤0.05 and ** p≤0.01. Error bars are standard error of the mean.

**Figure 10.**
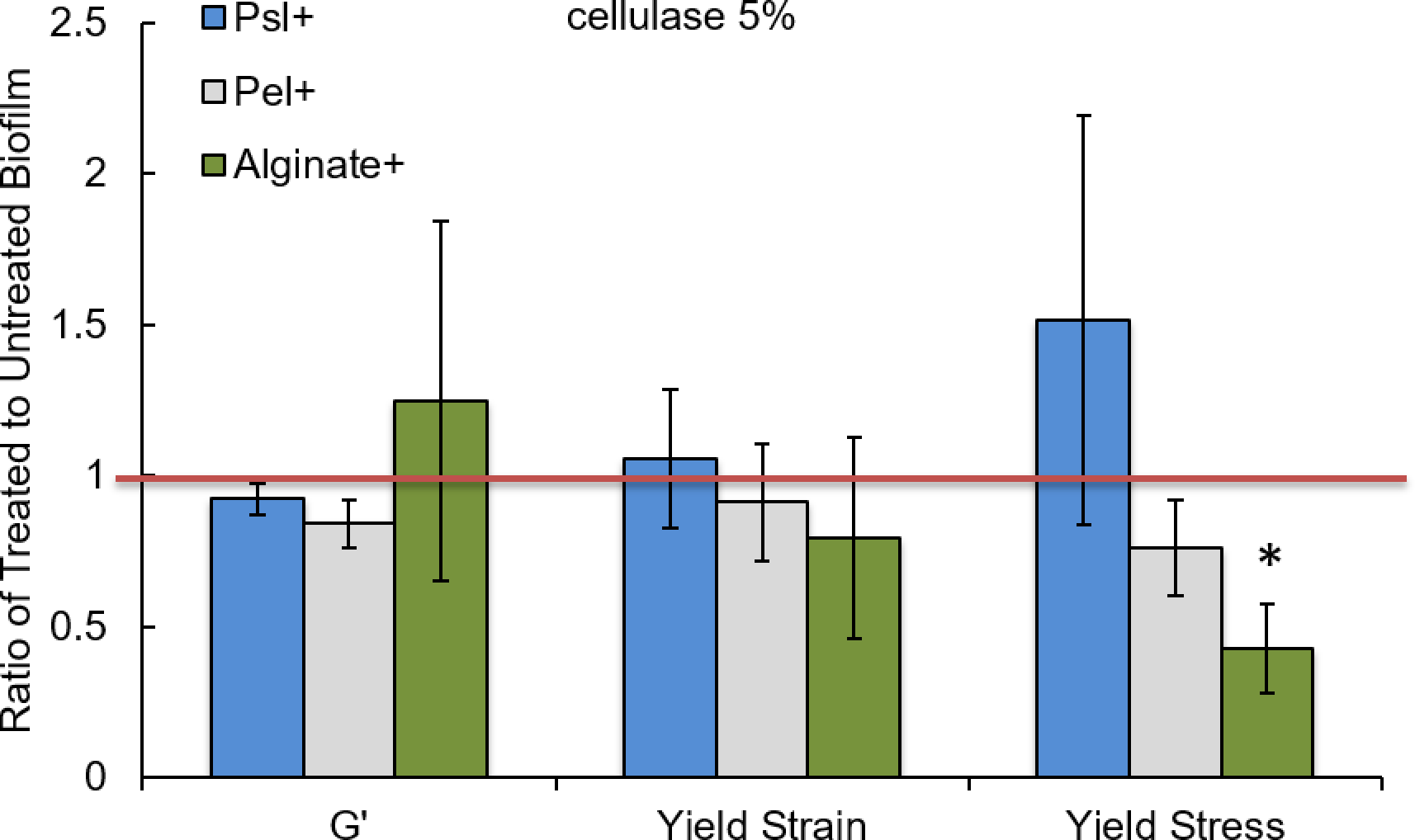
Treatment by 5% cellulase has minimal effects on biofilm mechanical properties. An alginate-dominant biofilm experiences a decrease in yield stress. * p ≤0.05 and ** p≤0.01. Error bars are standard error of the mean.

Furthermore, when we increase the dose of glycoside hydrolases used to treat WT biofilm from 5% to 10% to 20%, we do not see any statistically-significant trends (Figure S6 and S7). It may be that the biofilm compromise and dispersal previously seen with these hydrolases is not associated with significant changes in bulk biofilm mechanics and/or that it is not measurable by our rheometric technique. Our rheological studies are not sensitive to changes in adhesive forces (as long as the biofilm remains adhered to the rheometer tool, which it did in all cases), and the hour of treatment time may not be enough to fully capture the effects of enzymes on biofilms. If there are metabolic responses to the presence of enzymes, our assay is also unlikely to capture these changes—as we can see with there being no mechanical response to arginine treatment of biofilms, shown to be a disruptor via metabolic action by others.^31^ In addition, some environmental conditions may enhance or diminish the action of these hydrolases; indeed, cellulase has been shown to be pH-sensitive, with greater biofilm inhibition at pH 5 than pH 7.^37^

### Enzymatic treatment of biofilms grown in vivo

To assess the degree to which the *in vitro* results described above may provide a guide to the treatment of biofilm infections *in vivo*, we tested alginate lyase and DNase I against biofilms grown in a mouse model of chronic wound infection. For this, we did not measure biofilm mechanics but rather biofilm dispersal, because we have previously shown that biofilm dispersal in combination with antibiotic therapy is an effective treatment for biofilm infections.^8, 47^ Dispersal happens when the bacteria in a biofilm infection convert to the planktonic state; we anticipated that compromising biofilm mechanics would promote dispersal.

We used bacterial strains that form Alg+, Psl+, and Pel+ biofilms *in vitro*; it is worth noting that we have not independently characterized the EPS composition of the corresponding biofilm matrices for biofilms grown *in vivo*. For nominally-Psl+ biofilms, there was 48% greater dispersal when treated with a mixture of alginate lyase and DNase than when treated with a control (Figure 11). For nominally-Alg+ biofilms, there was only a 16% greater dispersal when treated with the enzyme mixture than when treated with a control; for nominally-Pel+ biofilms, there was only a 3% greater dispersal when treated with the enzyme mixture than when treated with a control.

**Figure 11.**
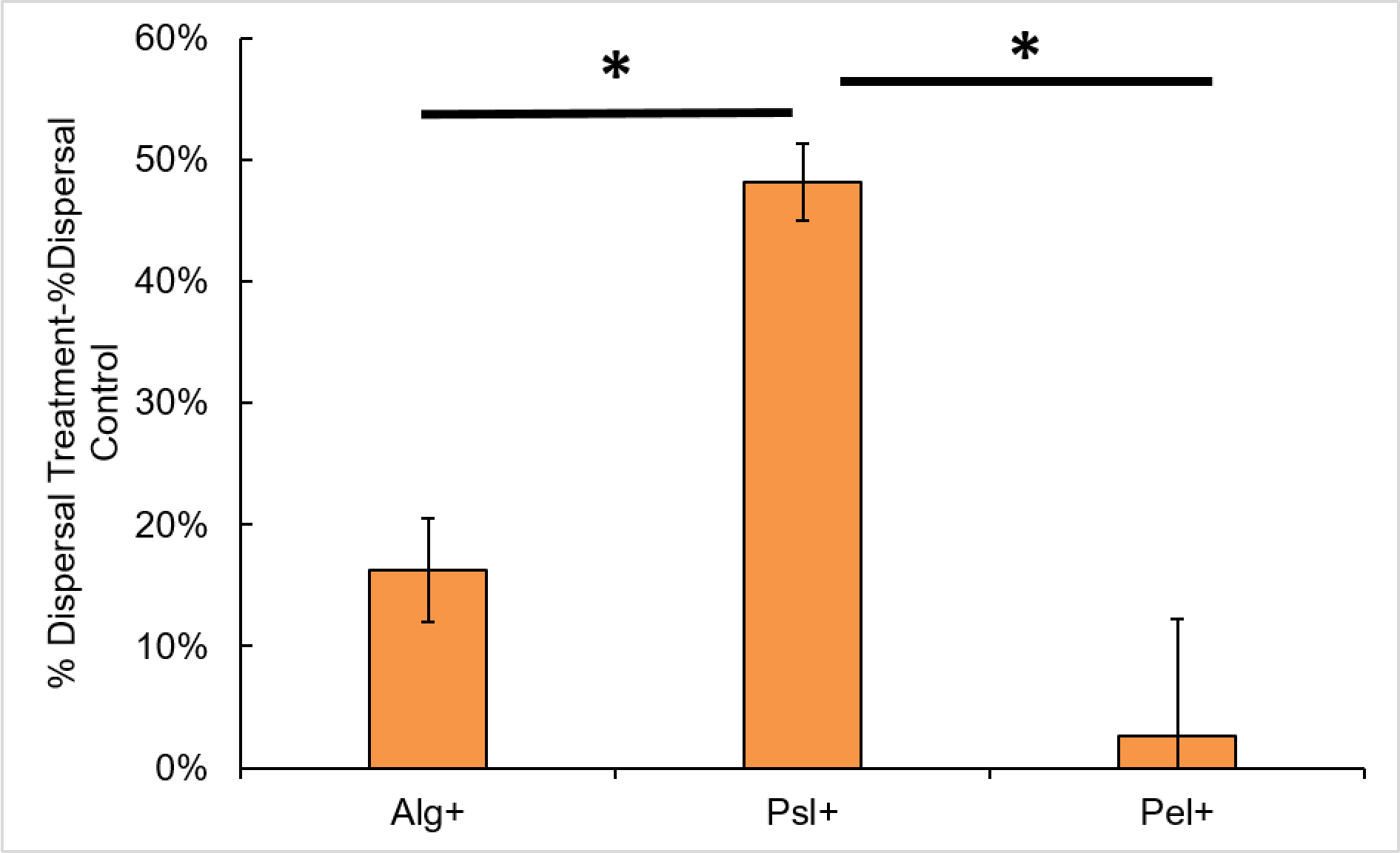
A mixture of alginate lyase and DNase elicits the greatest dispersal from Psl-dominant biofilms. Data shown are the dispersal measured for the treated sample minus dispersal measured for vehicle control. When 3-day biofilms grown in the mouse chronic wound model are treated *ex vivo* with 200 U alginate lyase and 500 U DNase, Psl-dominant biofilms experience greater dispersal, compared with the vehicle control, than either alginate-dominant or Pel-dominant biofilms. * p ≤0.05. Error bars are standard error of the mean.

These findings do not parallel our mechanical results for biofilms grown *in vitro*, for which the greatest response was seen for Alg+ biofilms treated with alginate lyase and Pel+ biofilms treated with DNase I. Furthermore, we previously found that the generic glycoside hydrolases, which have only minimal effect on biofilms grown *in vitro*, result in far more dispersal on biofilms grown in the same manner *in vivo* than do the specific enzymes.^8, 47^ One explanation for this apparent discrepancy might be that the changes in biofilms associated with dispersal are not primarily mechanical in nature. Another possibility is that the matrix produced *in vivo* might not have the same population of polymers as that produced by the same strain *in vitro*, and therefore might not have appropriately specific targets for the alginate lyase and DNase I.

## Discussion

### Summary of Results

For biofilms grown *in vitro*, alginate lyase and DNase I cleave alginate and eDNA polymers, respectively. For biofilms in which the target polymer is an important contributor to mechanics, this decreases yield strain and increases the diffusive transport of water through the biofilm matrix; we expect that this should correlate with increased diffusive transport of antibiotics as well. The increase in elastic modulus that we find upon cleaving a dominant polymer type results from drying due to faster water transport, and is therefore not a direct effect of cleaving the mechanically-dominant polymer; a biofilm surrounded by a fluid or high-humidity environment, as it would be in the body, would not dry and therefore would not experience an increase in elastic modulus. The dramatic effect of DNase I on Pel-dominant biofilms but not Psl-dominant or alginate-dominant biofilms indicates that extracellular DNA is a dominant mechanical component of the Pel polymer network, consistent with others’ finding that Pel associates with eDNA in the biofilm matrix.^20^

These results are not paralleled when we measure the effects of enzymatic treatment on the dispersal of biofilms grown *in vivo*. This apparent discrepancy both signals that dispersal may depend more strongly on non-mechanical properties than on mechanical properties and highlights the importance of considering growth environment when developing strategies for biofilm treatment. Growth in a living host could lead to changes in polysaccharide production through changes in gene expression and to other changes in matrix content through the incorporation of host material.^25, 48^ Understanding how biofilm dispersal happens if biofilm mechanics is not an important contributor, and how the content and enzymatic vulnerabilities of biofilm matrices depend on *in vivo* growth conditions, both seem like worthwhile avenues of future research based on the findings we present here.

### Enzymes needed to compromise Psl matrix

We have previously shown that Psl is important for biofilm stiffness.^21^ In this study, we find that for Psl-dominated biofilms grown *in vitro*, every enzyme used in this study showed no effect on mechanics and at most a minor effect on matrix structure (Figure 1-2,S4-S5). Psl may be particularly difficult to mechanically compromise. It has recently been shown that when Psl complexes with the protein CdrA, the biofilm is protected from proteolysis.^49^ We found previously that CdrA is required for the mechanical robustness of the Psl polymer network.^21^ Thus, the interaction of Psl and CdrA both enhances biofilm mechanics and protects the matrix from degradation. CdrA binds to mannose groups on Psl, and we and others have seen that when soluble mannose monomers are added to a liquid culture of Psl-producing bacteria, they act as a competitive binder for Psl and the bacteria fail to aggregate (Figure S8).^50^ However, we were not able to compromise the mechanical properties of the matrix by the addition of mannose after the biofilm was mature.

### Interpretation of Rheological measurements

Interpreting rheological measurements requires careful analysis and, often, additional experimentation, exemplified in the work we present here. The increase in elastic modulus seen after targeted treatment by the alginate lyase and DNase I enzymes may initially appear incongruous with the multitudes of evidence that biofilm networks are compromised by the addition of these enzymes.^12-14, 17-18, 32-33, 35^ However, due to a secondary effect of the enzymes—the biofilm drying—this result is actually another piece of evidence for matrix compromise.

### Future Directions: Other measurement methods

Our rheological measurements primarily measure cohesive properties of the matrix, being that the biofilms always remain fully adhered to the rheometer tool. Other measurement methods should be used to complete the mechanical picture of these polymer networks and reinforce rheological measurements. The mechanical contributions of individual polysaccharides to adhesive forces have yet to be quantified, although it is known that Psl contributes to permanent biofilm attachment, both Psl and Pel can play a role in initial attachment,^51–55^ and alginate is not required for surface attachment.^56^ In addition, how individual polysaccharides contribute to properties such as wetting behavior and hydration of the biofilm network, and how degradation of specific matrix polymers could give rise to alterations in the diffusion of water, antibiotics, and nutrients is still understudied.

## Acknowledgements

This work was supported by grants from the Cystic Fibrosis Foundation (GORDON1710), the National Institutes of Health (1 R01 AI121500-01A1 to VDG) (1 R21 AI137462-01A1 to KPR), the Ted Nash Long Life Foundation to KPR, and the National Science Foundation (1727544) to VDG. Bacterial strains used were gifts from Professor Steve Diggle (then, University of Nottingham; now, Georgia Institute of Technology) and Dr. Yasuhiko Irie (then, University of Nottingham; now, University of Dayton).

## Methods

### Bacterial Strains and Growth Conditions

The bacterial strains used for rheological studies and SEM are all in the PA01 background: wild-type (WT), Δ*wspF* Δ*pel* (which over-expresses Psl), Δ*wspF* Δ*psl* (which over-expresses Pel), and Δ*mucA* (which over-expresses alginate). Each of these strains constitutively expresses green fluorescent protein, so that any future experiments with fluorescent microscopy may be done with the same strain of bacteria. We grew the bacteria shaking in 4mL luria broth (LB) liquid media overnight at 37°C.

### Application of Biofilm Treatments for Rheology

To grow enough biofilm for rheological study, we spread 250μL of overnight growth on LB agar plates of standard size, 100mm × 15mm, and let these grow overnight at 37°C. Once the biofilms have grown overnight on LB agar plates, we apply our biofilm treatments by adding a liquid layer of treatment solution, 50μL to 100μL in volume, to the top of the lawn of biofilm on the plate. We use an L-spreader to gently—with minimal disruption to the biofilm—spread the liquid evenly over the biofilm. The treatments used in this experiment are Optizyme DNase I (Fisher), alginate lyase (Sigma), α-amylase from *Bacillus subtilis* (MP Biomedicals), cellulase from *Aspergillis niger* (MP Biomedicals), L-arginine hydrochloride (Fisher), and Salmon Sperm DNA (Fisher). For each treatment, we also treat with the solvent of the treatment material as a control. For example, the DNase I enzyme is buffered in 100mM Tris-HCl (7.5pH), 25mM MgCl_2_, and 1mM CaCl_2_, so the control for the DNase I treatments is this buffer solution alone. The treatment and control is then left to sit upright for an hour at 37°C. After an hour, the biofilm plate is ready for rheological measurement. Concentrations for the enzyme treatments are reported in activity units (U) per milliliter, where 1 U is the amount of enzyme that catalyzes one micro-mole of substrate per minute.

### Rheological Measurement

Rheology is done similarly as previously described in Kovach *et al*., with minor geometry changes.^21^ For this study, we used a stress-controlled AR 2000ex rheometer with a parallel-plate geometry with 8mm steel head. The biofilm is gently scraped from the surface of the agar onto the bottom plate of the rheometer; this typically only takes one plate of biofilm to fill the gap. The rheometer head is then lowered to a 500μm gap. Excess biofilm was trimmed to appropriately fill the gap; this takes approximately 60μL of biofilm. The rheological measurements were run at room temperature with the same solvent trap from Kovach *et al.* to stop evaporation of water from the biofilm while it was on the rheometer. ^21^

For each sample, we first run a frequency sweep from 0.1 to 200 rad/s at 1% strain to test the frequency dependence of the biofilm mechanics. In general, the biofilms have low frequency dependence from approximately 0.1 to 100 rad/s. After the frequency sweep, we run a strain sweep at 3.14 rad/s from 0.1 to 1000% strain. To determine the plateau G’, we pick a point in the middle of the plateau region of the strain sweep, generally at either 0.5% or 1% strain, and keep this choice constant for analysis of each biofilm treatment and its controls.

### Analysis of Rheological Data

Elastic modulus and yield strain were determined from the strain sweep data. Elastic modulus is determined to be the value of G’ in the plateau region of the strain sweep. To determine yield strain, we fit the plateau region to a linear fit and fit the “broken” nonlinear region to a power law. We report the intersection of these fits as the yield strain. The yield stress corresponding to this yield strain is then taken from the raw rheological data.

### Biofilm Drying Measurements

We grow biofilms as we do for rheology. We spread 250μL from liquid overnight onto LB agar plates of standard size, 100mm × 15mm, and let these grow overnight at 37°C. The biofilm is then gently scraped from the plate onto a weigh boat. For Δ*mucA* (Alg+) biofilms, we add 100μL of treatment such that the concentration is 200 U/mL alginate lyase (Sigma) in deionized water and for Δ*wspF* Δ*psl* (Pel+) biofilms, we add 100μL of treatment such that the concentration is 500 U/mL deoxyribonuclease I from bovine pancreas (Sigma) in 0.15M NaCl. The weigh boat with biofilm and treatment is then weighed. The biofilm is moved from the weigh boat onto a clean LB agar plate, as a sink for the excess water in the biofilm. This plate is then moved to an incubator at 37°C for one hour. After the hour, the biofilm is returned to its weigh boat and weighed. The alteration in weight is then calculated as water loss by 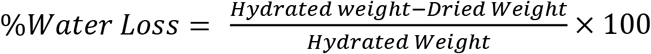.

### Statistical Significance

P-values are calculated using the Student one-tail t-test. In bar graphs, each value is tested against the null hypothesis that the reported ratio value is unity. P-values are reported as * p ≤0.05 and ** p≤0.01. Error bars are standard error of the mean.

### Scanning Electron Microscopy (SEM)

To image biofilms in SEM, we grow the biofilms on small glass pieces, 6mm × 1cm, which have been cut from standard microscope slides. We place these glass pieces into the wells of a 24 well plate. For growth of biofilm, we add 10μL of overnight growth to 1mL of LB in the wells. The biofilm is left to grow on the glass pieces for 24 hours as a static culture. At 24 hours, we gently pull the supernatant from the wells, leaving the biofilm growth on the glass pieces intact, and add the enzyme treatment to the biofilm. We add 500μL of treatment and controls to the wells. For SEM, we use alginate lyase (Sigma) in deionized water and deoxyribonuclease I from bovine pancreas (Sigma) in 0.15M NaCl. The alginate lyase treatment is at a concentration of 200U/mL and the DNase I treatment is 500U/mL. The treatment and the controls are let to sit for one hour at 37°C. After one hour, we wash the treatment out gently with PBS twice.

For standard fixation steps for SEM, we then move the glass pieces into a new 24 well plate. The first step is fixation with 1mL 4% glutaraldehyde and 2% paraformaldehyde in 0.1M cacodylate buffer and 2mM Ca^2+^ and 4mM Mg^2+^; this step causes proteins in the sample to irreversibly cross-link. The samples are left in the aldehyde solution overnight at room temperature. After overnight aldehyde fixation, the aldehyde is washed from the samples with 0.1M cacodylate buffer 3 times with 10 minutes between each wash. We then stain the samples with a mix of 4% osmium tetroxide and 4% potassium ferrocyanide (OsFeCN) at 1:1 ratio, giving us a solution of reduced osmium; the staining is set by microwaving at vacuum at 100W for 2 minutes twice, with 2 minutes of wait time between. Reduced osmium crosslinks with the lipids in the membrane of the bacterial cells, increasing membrane contrast to electrons for imaging. After the osmium fixation, we wash the osmium solution from the wells of the plate with deionized water. Once the osmium solution has been removed, we dry the sample.

The first step in drying is replacing the water in the sample with ethanol by placing the samples in 50%, 75%, and 95% ethanol for 10 minutes each sequentially. Then the sample sits in 100% ethanol for 10 minutes twice. We then dry the samples with a critical point drier. Once the sample is dried, it is fixed to SEM mounts using carbon tape and grounded with colloidal graphite paint around the edges of the sample. We then sputter coat the sample with 14nm platinum/palladium. For imaging, we use a ZeissSupra40 Scanning Electron Microscope operated at 5keV. The detector used captures type II secondary electrons. These electrons scatter at a wide angle and therefore capture compositional and some topographical information of the surface due to larger penetration depth, as opposed to type I secondary electrons that scatter at a smaller angle and are more sensitive to topological information.

### Murine Chronic Wound Model

Strains of *P. aeruginosa* were grown in baffled Erlenmeyer flasks at 200rpm in LB at 37°C, from which planktonic cells were harvested for injection into the wound. A full description of the chronic wound model used in this study can be found in previous work.^9, 57-60^ Briefly, mice were anesthetized by intraperitoneal injection of sodium pentobarbital. After a surgical plane of anesthesia was reached, the backs were shaved and administered a full-thickness, dorsal, 1.5 × 1.5 cm excisional skin wound to the level of panniculus muscle with surgical scissors. Wounds were then covered with a semipermeable polyurethane dressing (OPSITE dressing; Smith & Nephew®), under which 10^4^ bacterial cells were injected into the wound-bed. Biofilm formation was allowed to proceed for 72 hours, after which the mice were euthanized, and the wound-beds were harvested for *ex vivo* treatment with vehicle control, Alginate Lyase + DNAse, or Alpha-Amylase + Cellulase. Colony forming units (CFUs) were determined via serial dilution plating on *Pseudomonas* isolation agar, and percent bacterial cell dispersal was calculated by finding the quotient of the total CFU (biofilm-associated plus planktonic) divided by the planktonic CFUs (in the supernatant). Animals were treated humanely and in accordance with protocol #07044 approved by the Institutional Animal Care and Use Committee at Texas Tech University Health Sciences Center in Lubbock, Texas.

---

**Figure S1.**
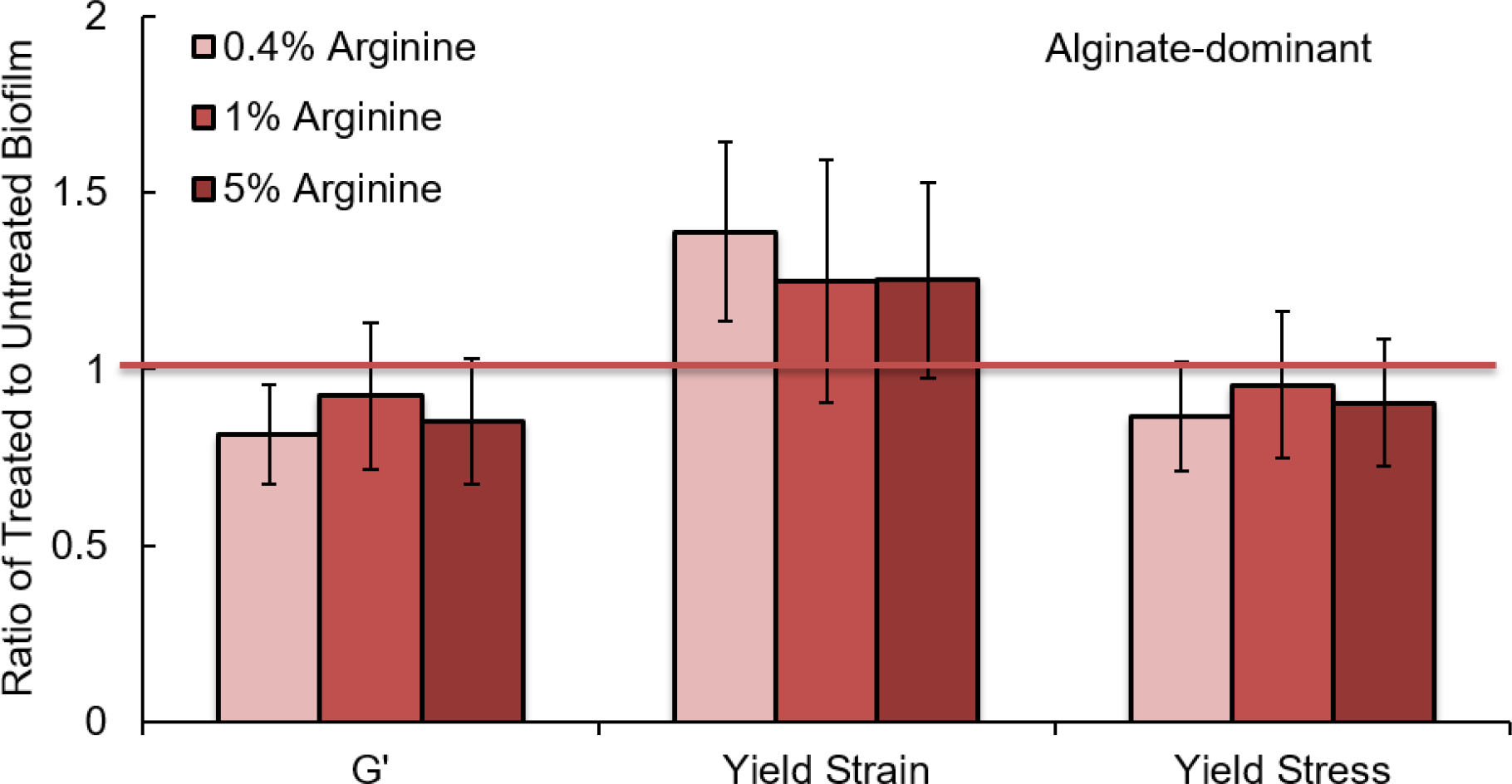
Arginine does not trigger mechanical changes in an alginate-dominant biofilm. Arginine is an amino acid constituent of alginate lyase. * p ≤0.05 and ** p≤0.01. Error bars are standard error of the mean.

**Figure S2.**
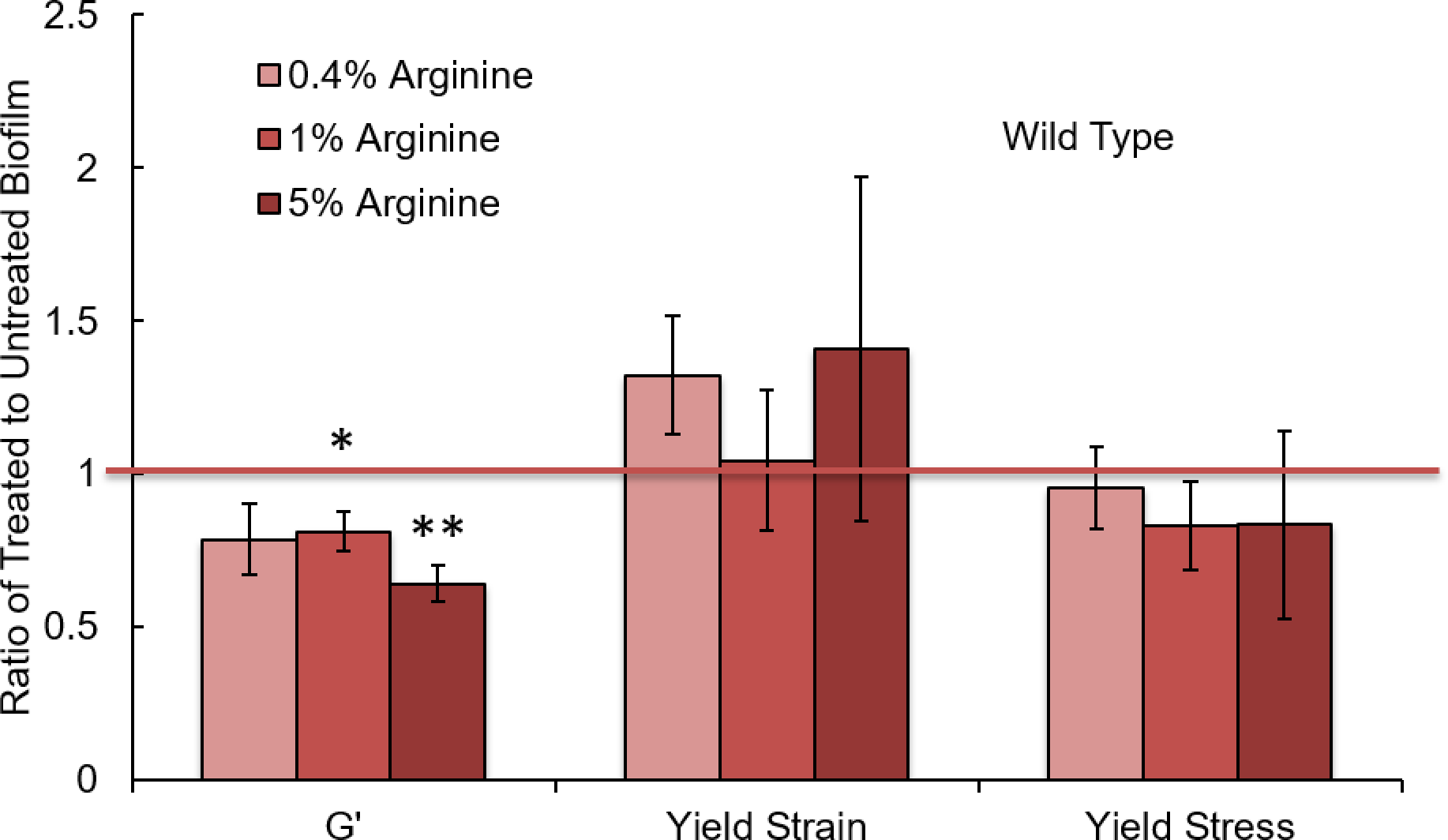
Increasing arginine concentration softens wild type biofilm. In a wild type biofilm, whose mechanical properties are dominated by Pel and Psl, the elastic modulus decreases at 1% and 5% arginine. In alginate lyase treatment, elastic modulus increased, but these treatments elastic modulus decreased. Therefore these mechanical changes are not like those found in alginate lyase treatment, and so unlikely to be correlated. * p ≤0.05 and ** p≤0.01. Error bars are standard error of the mean.

**Figure S3.**
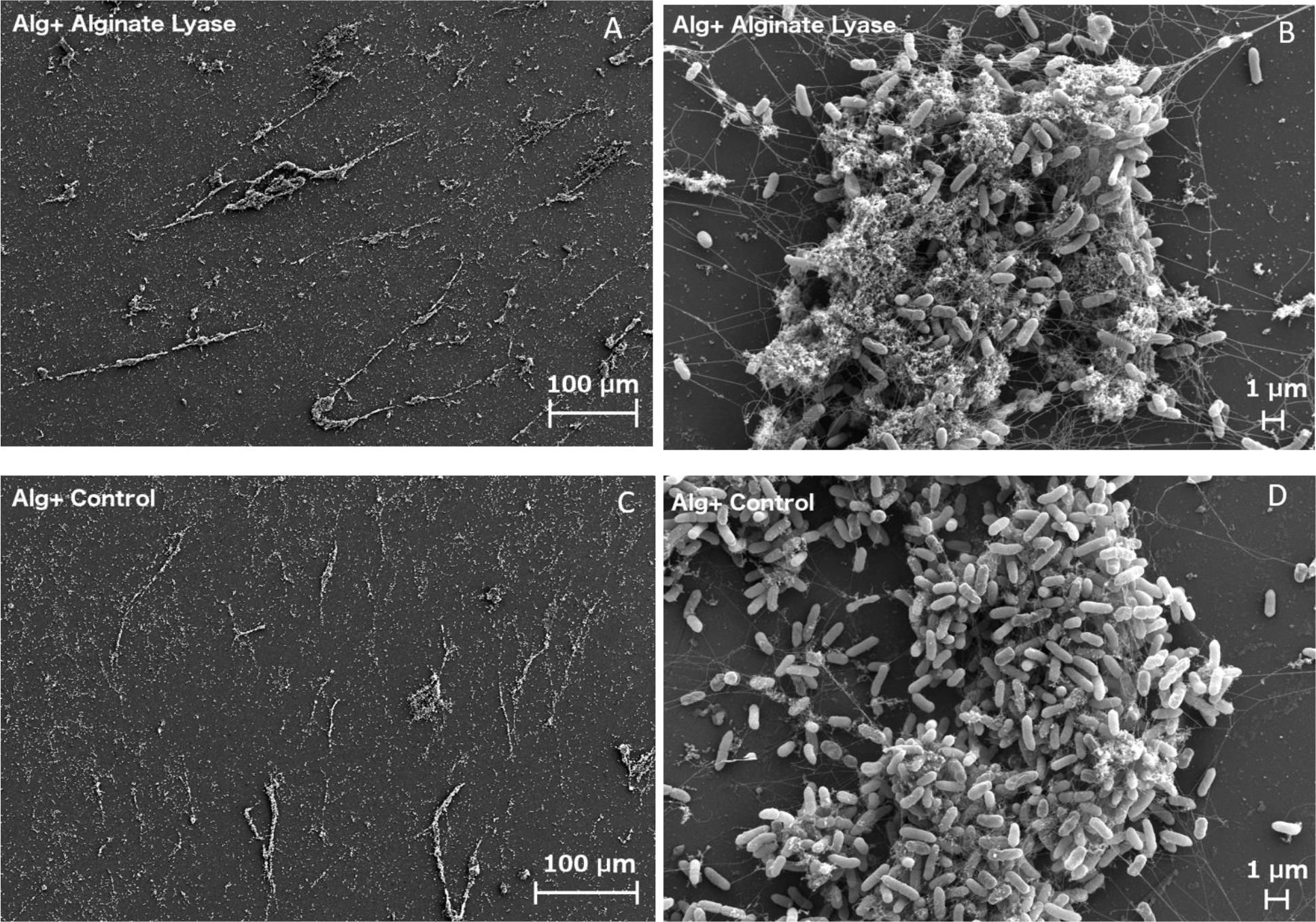
SEM images of alginate-dominant biofilms. Despite mechanical changes measured upon alginate lyase –treatment of alginate-dominant biofilms, the SEM images do not have striking visual differences. The long-range connectivity and structure of (A) treated and (C) control biofilms appears similar. The enzyme-treated biofilm (B) actually appears denser in polysaccharides than does the untreated biofilm (D).

**Figure S4.**
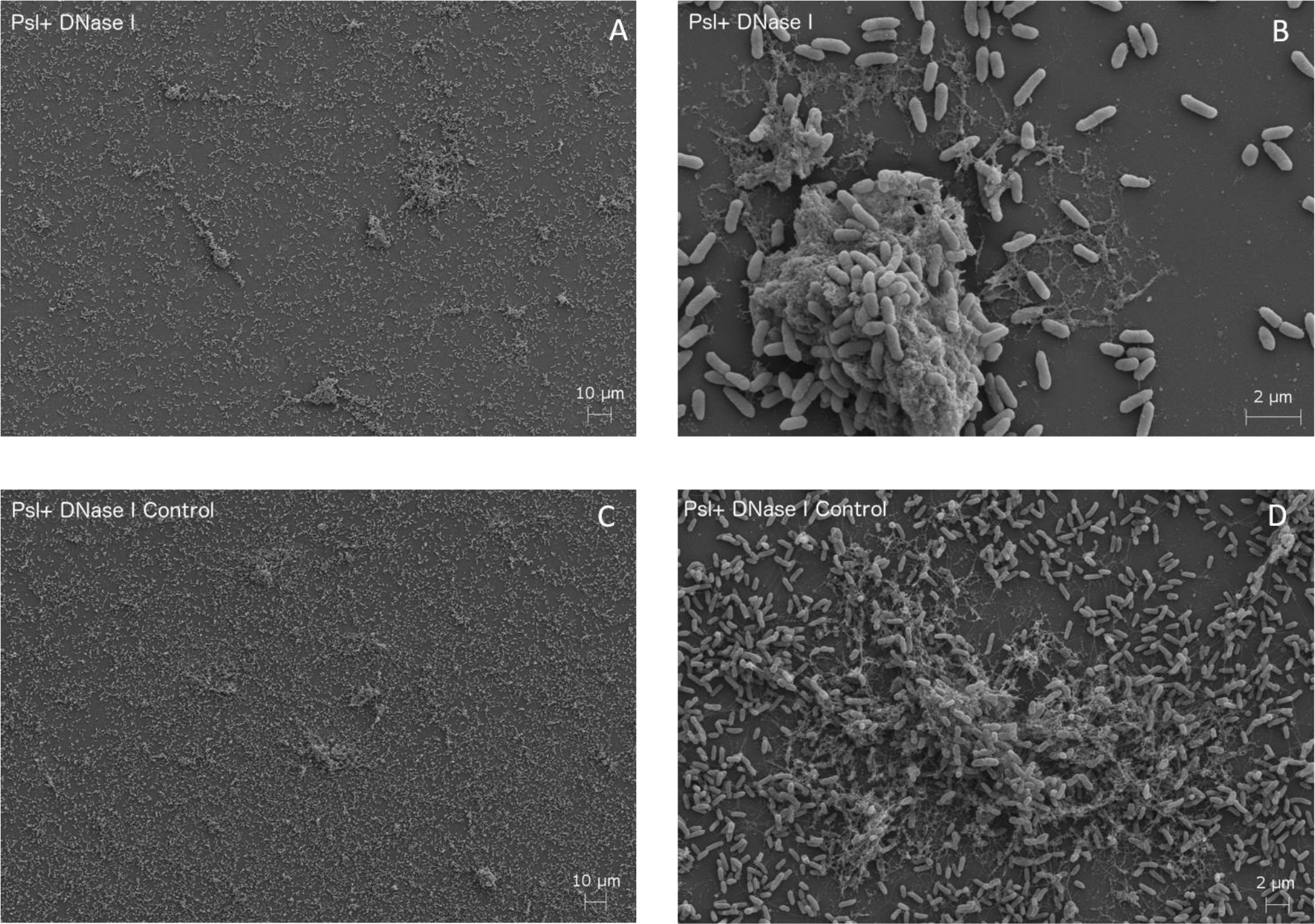
SEM images of Psl-dominant biofilms. At large scale, it is apparent that (A) biofilms treated by DNase I have fewer cells remaining on the surface than do (C) control biofilms. In addition, the DNase I-treated biofilm (B) appears to have a dense polysaccharide structure surrounding cells, while the untreated biofilm (D) has no dense structure, only polysaccharide strands present. This density change could be a result of dehydration due to enzyme treatment as seen in Pel-dominant biofilms treated by DNase I (Figure 6).

**Figure S5.**
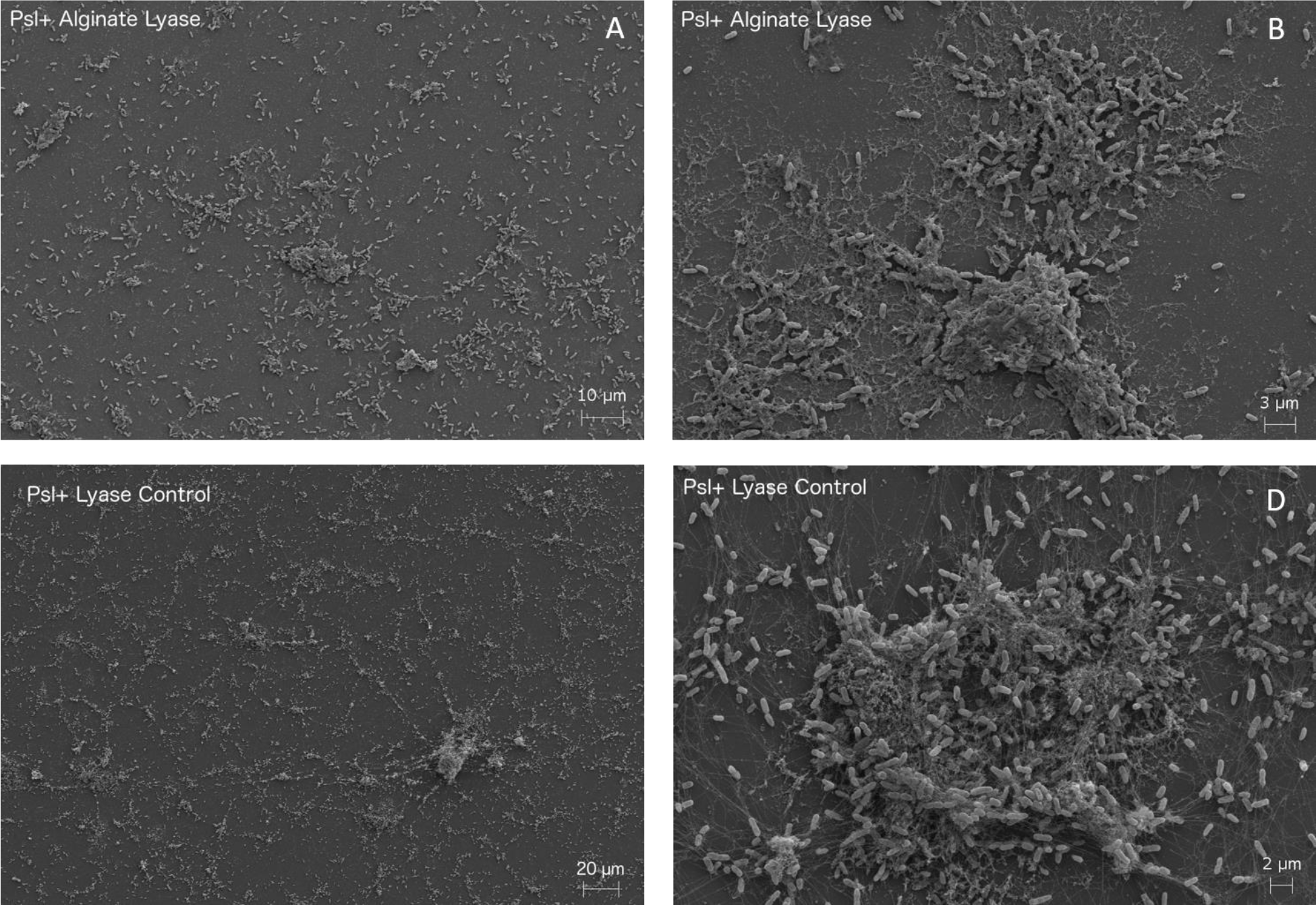
SEM image of Psl-dominant biofilms. (A) The biofilm treated with alginate lyase appears to have fewer cells remaining on the surface than does (C) the control biofilm, although this difference is not as striking as that seen in Figure S4 when the same type of biofilm is treated with DNase I. A stronger parallel with the effects of DNase I treatment is seen inthe dense polysaccharide structure in (B) biofilm treated with alginate lyase (compare with Figure S4b), and the less-dense polysaccharides seen in (D) the control biofilm (compare with Figure S4d). This density change could be a result of dehydration due to enzyme treatment as seen in alginate-dominant biofilms treated by alginate lyase (Figure 6).

**Figure S6.**
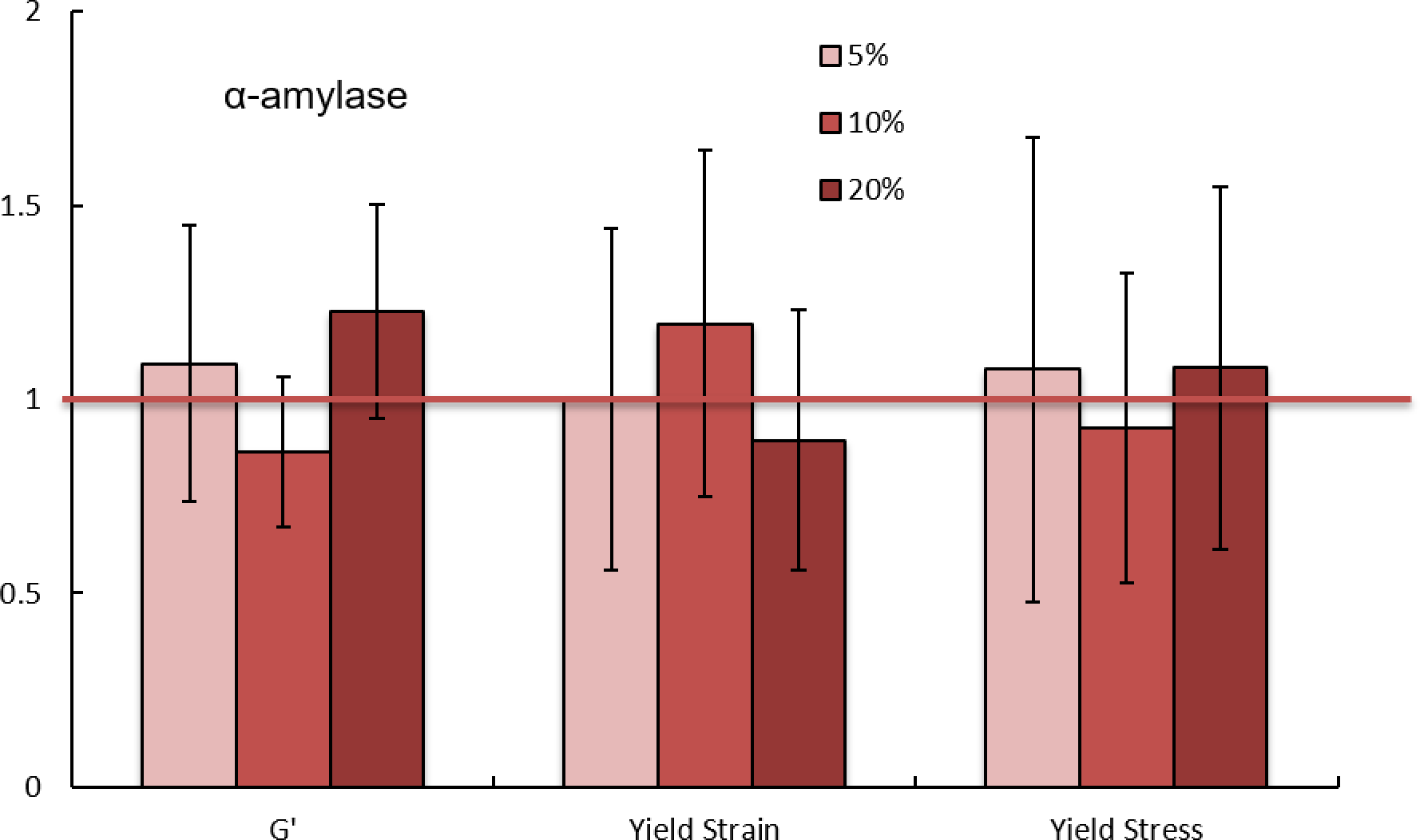
Increasing dosage of α-amylase has no effect on the mechanical properties of wildtype biofilms. The enzyme α-amylase has no effect on biofilm mechanical properties measured by bulk rheological methods. * p ≤0.05 and ** p≤0.01. Error bars are standard error of the mean.

**Figure S7.**
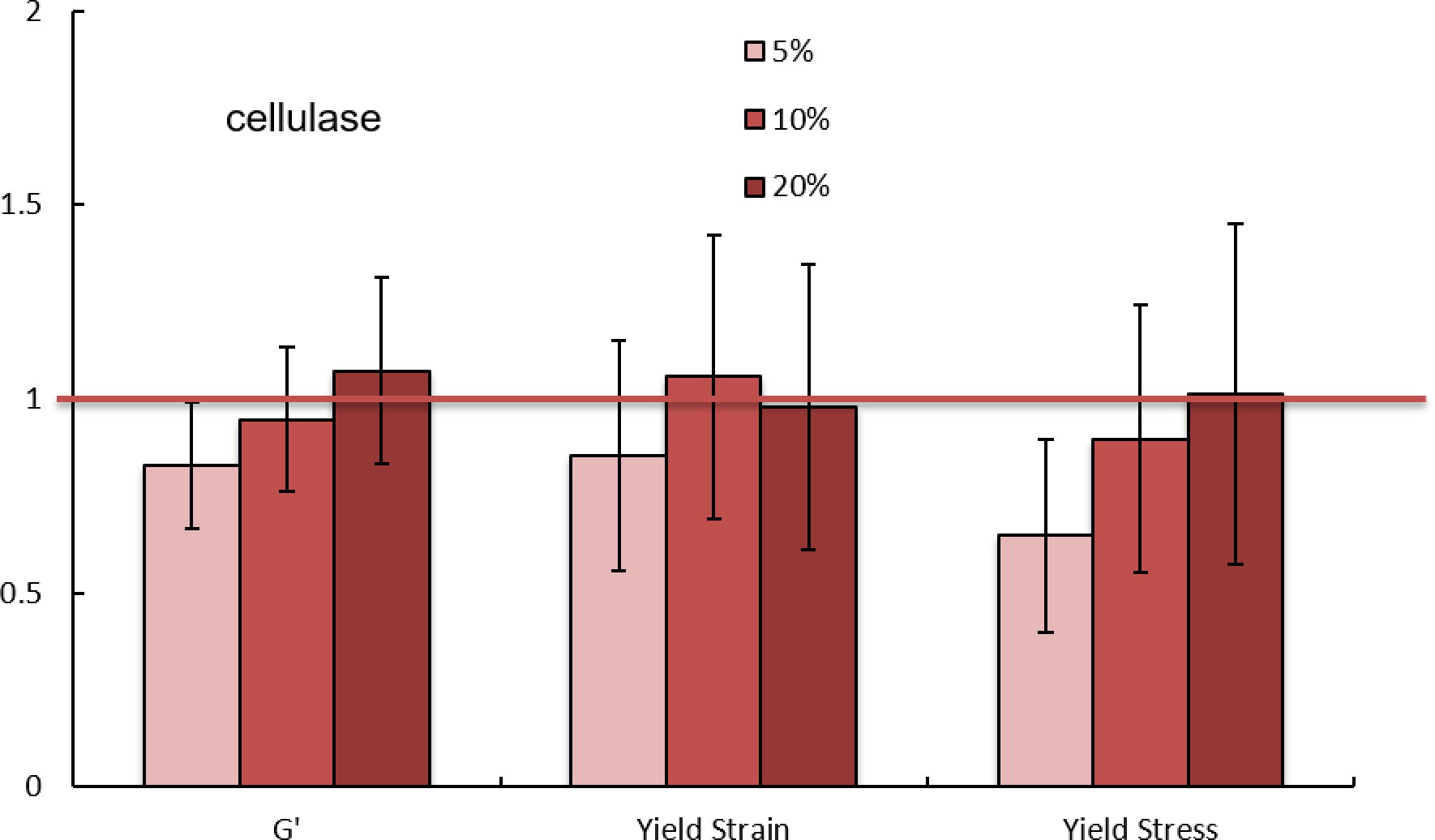
Increasing dosage of cellulase has no effect on the mechanical properties of wildtype biofilms. The enzyme cellulase has no effect on biofilm mechanical properties measured by bulk rheological methods. * p ≤0.05 and ** p≤0.01. Error bars are standard error of the mean.

